# Robust ciliary flows protect early Xenopus embryos from pathogens independent of multiciliated cell patterning

**DOI:** 10.64898/2025.12.19.694354

**Authors:** Athullya Baby, Alice Briole, Ayush Yadav, Isabelle Cheylan, Virginie Thomé, Camille Boutin, Umberto D’Ortona, Annie Viallat, Julien Favier, Etienne Loiseau, Laurent Kodjabachian

## Abstract

During the early stages of development, the skin of the *Xenopus* embryo is covered by around two thousand evenly distributed multiciliated cells (MCCs). This striking spatial distribution is believed to maximise the generation of superficial flows at the scale of the embryo. However, the specific role of regular MCC distribution and the physiological function of vigorous ciliary activity remain elusive. We investigate the extent to which superficial flows provide protection against external pathogens before the immune system matures, by combining experimental and computational approaches. First, we cultivated epithelial explants in order to quantify the distribution of MCCs, beating frequencies, and three-dimensional fluid flows. We then use these data to validate a computational fluid dynamics model. Using this model, seeded with structural data from whole embryos, our simulations reveal that the collective ciliary beatings create a robust liquid shield along the embryonic flank, which is highly effective at clearing pathogens from the vicinity of the epithelial surface. Through parametric analyses, we further demonstrate that this protective function is remarkably resilient. Indeed, the effectiveness of pathogen clearance is primarily governed by the overall characteristic velocity of the cilia and is less affected by moderate variations in MCC density and spatial organisation. Our findings suggest that, rather than optimising energy consumption, the biological system prioritises functional robustness to ensure reliable protection.

## Introduction

During the larval development of the amphibian *Xenopus laevis*, the skin is temporarily covered by approximately two thousand multiciliated cells (MCCs), which are evenly distributed amongst secretory cells that produce a protective mucus layer (1, 2). MCCs represent about 10% of the total number of cells on the surface of the embryo (3). The coordinated beating of their numerous cilia generates robust fluid flows directed from anterior-dorsal to posterior-ventral across the body surface (4, 5). MCCs emerge during embryonic development, and persist into the free-swimming tadpole stage, long after hatching, eventually disappearing at late Nieuwkoop–Faber stages (NF 40–47) through programmed cell death (apoptosis), or transdifferentiation into mucus-producing cells (2, 6–8). The precise physiological function of *Xenopus* MCCs and the vigorous flow they induce remains poorly understood. Several hypotheses have been proposed through works on different aquatic species, which include enhancement of respiratory gas exchange and, critically, clearance of pathogens in the vicinity of the embryo (2, 9–14). The hypothesis that cilia-powered flows protect embryos by preventing the attachment of post-hatching debris and pathogenic microorganisms is particularly compelling. This is supported by the observation that maximum ciliation occurs after hatching (the breaking of the vitelline membrane), and that peak ciliary activity coincides with a period when the immune system is still non-functional (2, 15, 16). In that respect, this clearance function would offer a dual safety mechanism alongside the biophysical barrier of the 6 µm-thick mucus layer covering the embryo’s surface (17).

The emergence of coherent cilia-powered flows depends on multiple parameters deployed at both cell and tissue-scales (5, 18–20). Such parameters, including cilia and MCC density, alignment, spacing and beating strength, are interdependent, making it difficult to evaluate experimentally their individual influence on flow production. Moreover, measuring a three-dimensional flow on the millimeter scale, especially over the non-transparent and non-symmetrical surface of a 3D embryo, is inherently difficult. While interesting theoretical and numerical approaches have been proposed, they often lack experimental validation at the tissue scale or comprehensive discussion regarding protection against infection (21).

A remarkable feature of *Xenopus* MCCs is their regular distribution across the embryonic skin, often at the nodes of a highly ordered network, that is established through active migration and homotypic repulsion, mediated by the Scf/Kit signalling system (22). Such a precise organization raises fundamental questions regarding its functional significance: is regular MCC spacing required for efficient surface cleaning, and does it represent an optimized configuration for pathogen clearance?

Here, we address these questions through a combined experimental and numerical approach. First, we establish a quantitative infection assay to directly assess the protective role of cilia-driven flows at the scale of the whole embryo. Second, we develop transparent and planar epithelial explants that enable to quantitatively characterize multiciliated cell organization, ciliary activity, and the resulting flow fields. These experimental measurements are then used to inform and validate a numerical fluid-dynamics model. Finally, we leverage this model to systematically and independently vary MCC beating strength, density, spatial organization, and alignment, and assess how each parameter shapes surface flows and governs pathogen clearance at the scale of the embryonic flank. This approach yields a quantitative description of MCC-driven flows across the epithelial surface. We show that MCC collectively generate a liquid shield that efficiently limits bacterial access to the embryonic surface. Remarkably, this protective barrier demonstrates high resilience to variations in both MCC density and spatial organization. This suggests that the system does not rely on fine MCC geometric optimization, but instead prioritizes robustness in physiological function, ensuring effective protection despite biological variability and structural imperfections in the ciliated epithelium.

## Results

### 1. Cilia-powered flows prevent infection of *Xenopus* embryos

Previous work suggested that long-term exposure (72 h) of *Xenopus laevis* embryos to *A. hydrophila* caused enhanced death, when cilia-powered flow was reduced (14, 17). To better quantify infection rates, we established a protocol to score the number of colony-forming units (cfu) from the lysate of individual embryos, exposed to *A. hydrophila* for 9 hours at stage 30, when the ciliated epidermis is fully functional (see Materials & Methods). Uninjected control embryos (Fig. 1A,B), as well as Cas9-injected control embryos (Fig. 1C,D), displayed normal ciliogenesis, powerful directional flow and showed very low infection rates (1/15 for either condition, average cfu=26 and 44, respectively). In contrast, embryos co-injected with Cas9 and a guide RNA targeting FoxJ1, a key regulator of motile ciliogenesis (23), lacked motile cilia, which abolished flow (Fig. 1E,F), as confirmed by bead velocity measurements (Fig. 1G). In this condition, the majority of embryos were infected (13/25, average cfu = 474), with approximately half of them dying during the 9 h exposure to *A. hydrophila* (6/13; Fig. 1H). This result confirms that cilia-powered flows help to protect *Xenopus* embryos from bacterial infection. Next, we wanted to evaluate the importance of regular MCC spacing for the resistance to infection by *A. hydrophila*. For this, we used axitinib, a chemical inhibitor of the Kit receptor, which was reported to alter MCC homotypic repulsion and spacing (22). Unexpectedly, we observed that exposure to axitinib upset MCC distribution, but also caused their premature disappearance (see Fig. S1). This compound phenotype prevented us from testing specifically the importance of MCC geometrical order for resistance against *A. hydrophila*. To overcome this experimental limitation, we developed a combined approach utilizing *in vitro* explants and a numerical model to facilitate the evaluation of how individual parameters of the system impact flow generation and pathogen clearance.

**Fig. 1.**
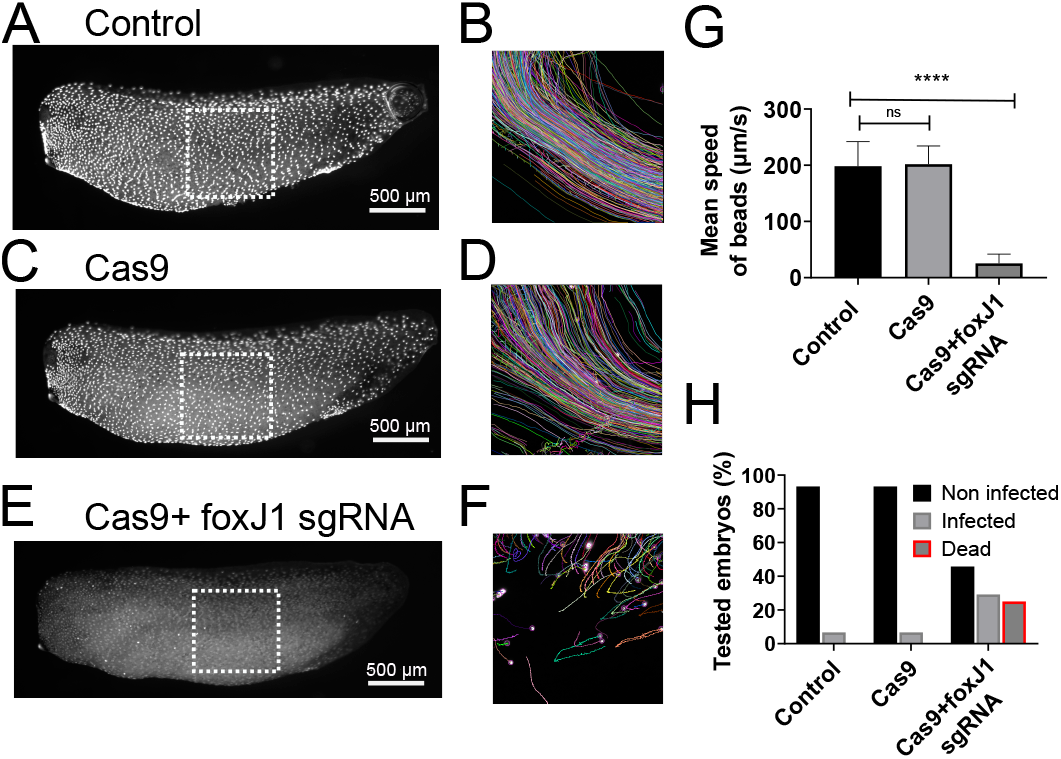
MCCs are required to limit infections of *Xenopus* embryos by *Aeromonas hydrophila*. Distributions of MCCs, visualized by immunofluorescent labeling of acetylated tubulin, on the flank of embryos, together with trajectories of fluorescent beads propelled by ciliary activity in control embryos (A) & (B) and for a Cas 9 injected embryo (C) & (D). For an embryo co-injected with Cas9 and guide RNA against FoxJ1, the main gene involved in ciliogenesis, cilia are absent (E) and beads are not transported (F). (G) Quantification of the velocity of beads above the flank of embryos.(H) Infectability of embryos by *Aeromonas hydrophila*. The absence of cilia due to foxJ1 knockout causes the lack of flow and enhanced infection rate.

### 2. Multiciliated cell organization and flow generation in embryos and explants

Here, we establish an experimental framework to quantify the spatial organization of MCCs and the flows they generate. Because direct three-dimensional flow measurements are not feasible on embryos due to their opacity and curved geometry, we use planar epidermis explants. We first compare static features of MCC organization in embryos and explants, and then quantify key dynamic parameters of ciliary activity and flow generation that will be used to parameterize the numerical model in the following section.

To experimentally perturb MCC spatial organization in explants, we use axitinib. While it induces premature MCC loss in embryos, it disrupts MCC spacing without affecting MCC survival or ciliary activity in explants. This difference is consistent with previous reports showing that MCC disappearance requires mesoderm-derived signals (7, 8).

#### Comparison of MCC spatial organization in embryos and explants

Embryos and explants were prepared at comparable developmental stages (around NF 30; see Materials and Methods). We compared MCC spatial order, density, and morphology between whole embryos and explants, under both control and axitinib-treated conditions.

Figure 2A shows the spatial distribution of MCCs on the flank of a fixed embryo. MCC positions were used to compute Delaunay triangulations (Fig. 2B) and derive a geometric order parameter quantifying MCC spatial organization (see Materials and Methods). Figures 2C and D show a representative planar epithelial explant and the corresponding MCC distribution obtained from live imaging. Untreated embryos and explants exhibited similar levels of spatial order (Fig. 2E), indicating that explant preparation preserves global MCC organization. In contrast, axitinib-treated explants displayed increased heterogeneity in MCC spacing, confirming that axitinib effectively perturbs MCC spatial organization. We then examined basic MCC morphological features. Individual MCC surface area was comparable between embryos and explants (Fig. 2F), indicating that explant culture largely preserves MCC morphology. In contrast, MCC density was significantly reduced in explants compared to embryos, reflecting tissue spreading on the substrate (Fig. 2G). Despite this reduction, MCC density in explants remained sufficiently high to generate robust cilia-driven flows.

**Fig. 2.**
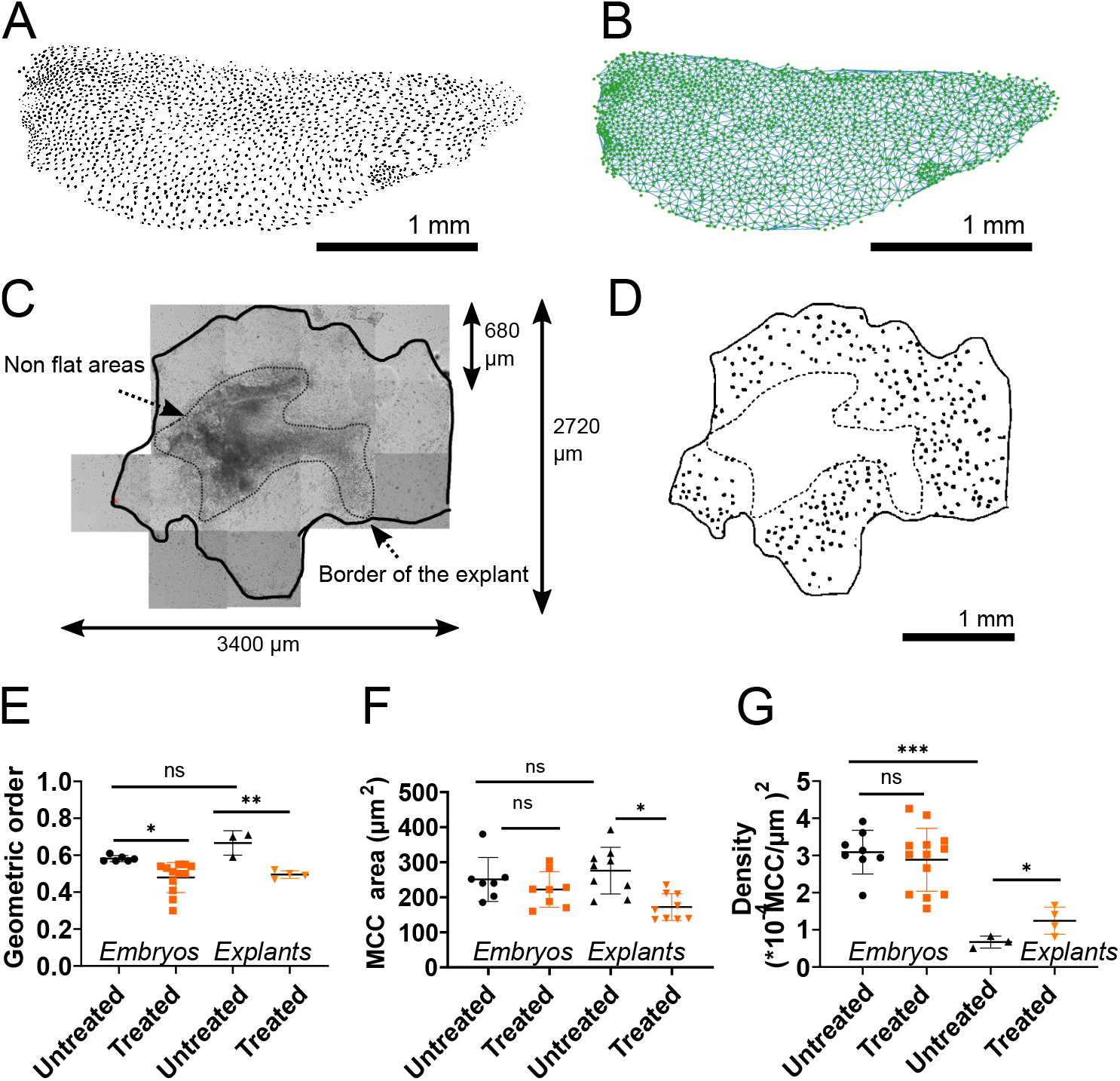
Static parameters of the ciliated epidermis. (A) Spatial distribution of the multiciliated cells (MCCs) on the flank of an embryo. Binarised image computed from a fluorescent image (see methods section). (B) Delaunay’s triangulation, computed from the image on panel (A), is used to compute the geometrical order. (C) Characteristic epidermal explant (3400 µm*2720 µm) visualized through stitching of 15 imaging tiles (680 µm*680 µm; see Materials & Methods). The region inside the dashed line corresponds to location where the tissue is not perfectly flat. (D) MCC distribution corresponding to the explant in (C). Binarised image computed from movie acquired in bright field (see methods). (E) Comparison of the geometrical order on embryos and on explants for both control and axitinib-treated conditions. (F) Comparison of the surface area of MCCs measured on fixed embryos and on live explants for both control and axitinib-treated condition. For explants, a correction is applied as described in the method section to compare with measurements done on fixed samples. (G) Comparison of the MCC density on embryos and explants for both control and axitinib-treated conditions.

Together, these results establish epithelial explants as a faithful proxy of the embryonic epidermis for studying MCC-driven flows. Moreover, axitinib treatment increases heterogeneity in MCC spacing while preserving a fully ciliated epithelium, enabling selective perturbation of spatial organization without affecting MCC viability.

#### Dynamic features of ciliary flow in explants

We performed a quantitative characterization of cilia-driven flows on explants by combining particle image velocimetry (PIV) of fluorescent microbeads with measurements of ciliary dynamics, including coordination and beat frequency, in both untreated and axitinib-treated conditions.

Ciliary beat directions were analyzed in both untreated and axitinib-treated explants (Fig. 3A,B), and the coordination was quantified by a polarity order parameter *P*, with *P* = 1 corresponding to perfectly aligned beat directions and *P* = 0 to random orientations (see Materials and Methods). Polarity order decreased gradually with increasing intercellular distance, reflecting a progressive loss of long-range correlation (Fig. 3C), which varies among explants. Notably, a strong degree of orientation, reaching up to 0.9, can be maintained at distances of up to 500 µm. Ciliary beat frequencies ranged from 8 to 16 Hz, with a mean value of approximately 13 Hz (Fig. 3D).

**Fig. 3.**
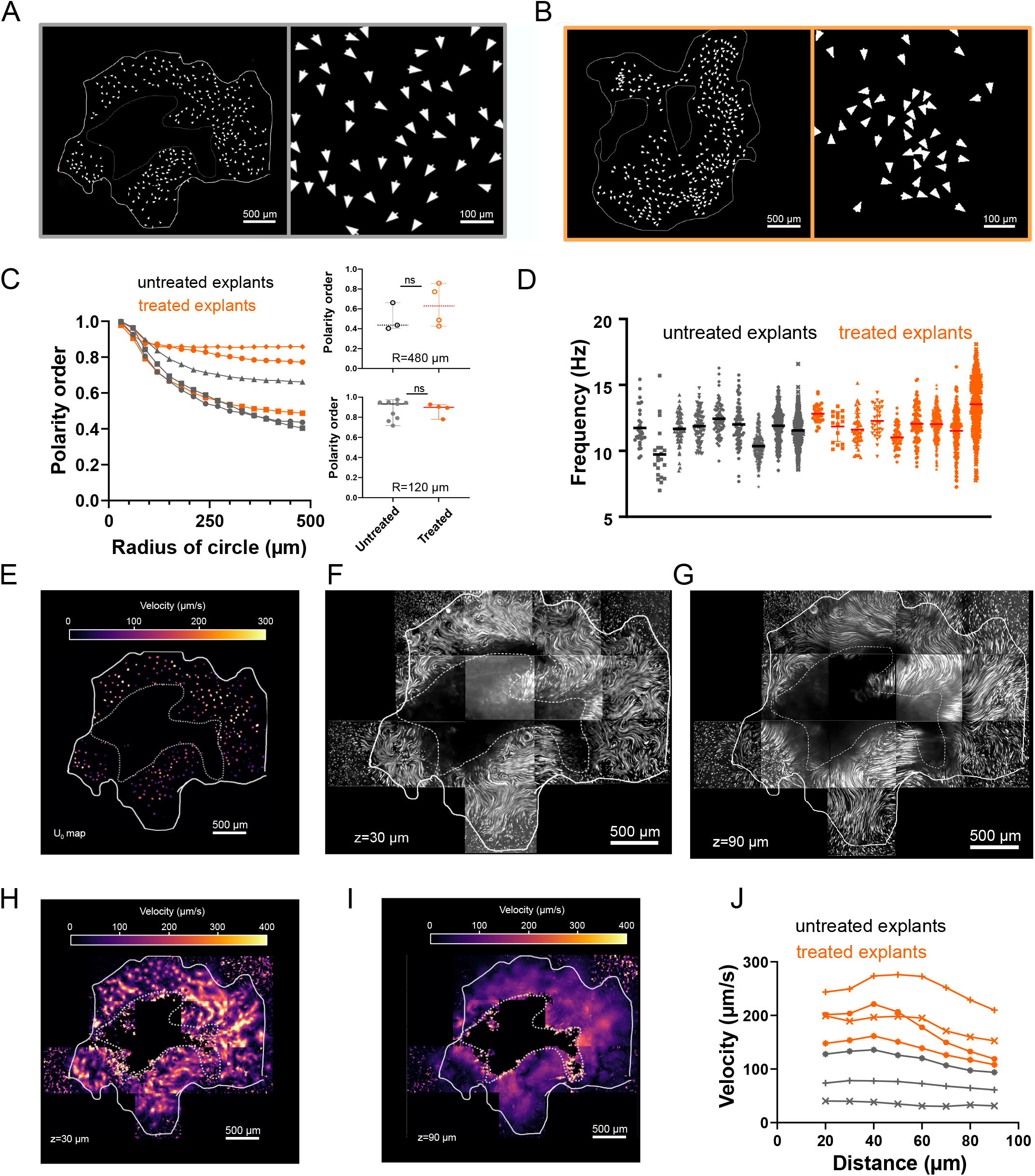
(A) & (B) Ciliary beat directions for a control explant (A) and an axitinib-treated explant (B). Right panels show magnified views to better visualize the coordination of beat directions. (C) Coordination of beat directions quantified by the polarity order over increasing length scales, from neighboring cells (≃ 50 *µ*m) up to 500 *µ*m. Long-range polarity analysis could only be performed on the largest explants. No significant difference is observed between control and axitinib-treated conditions. (D) Distributions of ciliary beat frequencies for both control and axitinib-treated explants. (E) *U*_0_ map corresponding to the velocity measured at the tip of the cilia for each multiciliated cell. (F) & (G) Visualization of the streamlines at two heights, *z* = 30 *µ*m and *z* = 90 *µ*m, above the surface of an explant. Streamlines are obtained by computing the standard deviation over 100 frames (≃ 1 s) of flowing fluorescent beads. (H) & (G) Maps of the flow velocity magnitude at different distances from the epithelial surface. (J) Average flow velocity profiles as a function of height above the epithelial surface for control and axitinib-treated explants. Profiles are shown for the largest explants analyzed, which allow reliable spatial averaging.

Local fluid forcing generated by multiciliated cells, denoted *U*_0_, was quantified by measuring the local velocity, using PIV, immediately above the tips of MCC cilia (Fig. 3E). The distribution of *U*_0_ exhibited substantial heterogeneity across individual cells within the same explant, likely reflecting variations in cilia number, and beat frequency.

No statistically significant differences were observed between untreated and axitinib-treated explants in ciliary co-ordination (Fig. 3C), beat frequency (Fig. 3D), or the overall distribution of local forcing amplitudes (Fig. S5).

The collective flow resulting from this local forcing was characterized by reconstructing streamlines from the PIV measurements at different heights above the epithelial surface (Fig. 3F,G). As the distance from the tissue surface increased, streamlines became progressively smoother and more regular. Bead trajectories were largely confined to the imaging plane, with occasional vertical displacements indicating the presence of three-dimensional flow components (Movie S2), which could not be quantified directly with our experimental setup.

Velocity maps showed heterogeneous and fast flows near the cilia tips, that became progressively smoother and more homogeneous farther from the epithelium (Fig. 3H,I). At the tissue scale, mean flow velocity profiles have similar shapes between untreated and axitinib-treated explants (Fig. 3J). However, the axitinib-treated explants exhibit higher flow amplitudes. The data shown in this figure correspond to the largest explants obtained experimentally, selected to allow robust statistical analysis over a wide field of view (untreated explants 7-9, see Table S3; axitinib-treated explants 6-9, see Table S4). The axitinib-treated ones also exhibit higher MCC density and larger local forcing amplitudes (*U*_0_), which are likely responsible for the increased flow amplitudes.

Overall, we show that axitinib treatment alters the spatial organization of multiciliated cells while leaving ciliary coordination, beat frequency, and local forcing largely unchanged. Variations in flow amplitude observed at the tissue scale likely arise from the combined influence of multiple parameters and cannot be disentangled experimentally, motivating the use of a numerical model to independently assess their respective roles in flow generation.

### 3. Numerical model of a ciliated epithelium

#### The model

Our numerical model, detailed in the Materials and Methods section employs a three-dimensional Lattice Boltzmann method to simulate the fluid dynamics above the embryonic tissue. The ciliated cells are incorporated using the Immersed Boundary Method (IBM), where each cell is represented as a point force acting on a Lagrangian point. These forces, which represent the net ciliary propulsion, are located at a height of 20 µm above the epithelial surface, corresponding to the approximate height of the cilia.

Figure 4A provides a schematic representation of MCCs and the *Xenopus* epithelium within our model. The input parameters for the model are the position, direction, and magnitude of these point forces representing the ciliated cells. The positions of the point forces directly correspond to the centroids of MCCs obtained from our experimental images (Fig. S3 & S4), their direction, in the horizontal plane, is determined based on the measured ciliary beating directions obtained from experimental data, and their magnitudes are directly obtained from the *U*_0_ values measured on our explant experiments.

**Fig. 4.**
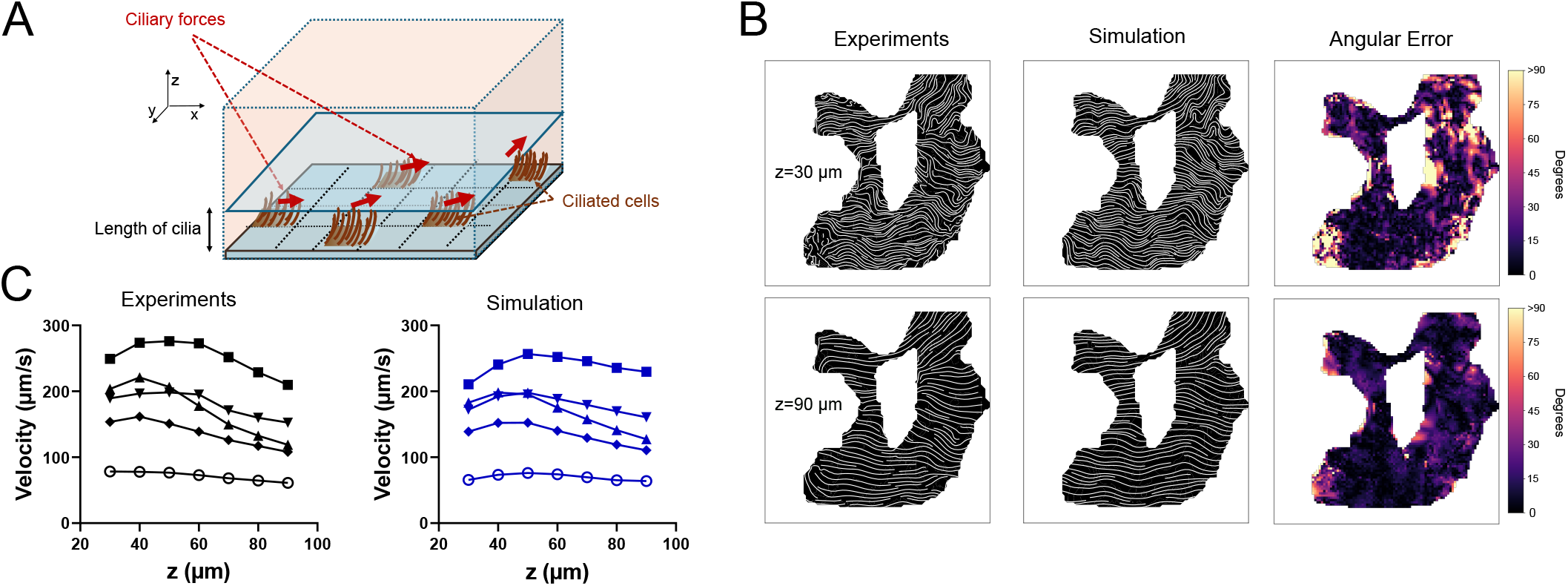
Validation of the numerical model for ciliated epithelia. (A) Schematic of the virtual epithelium. Ciliated cells are modeled by point forces. (B) Comparison between measured streamlines and streamlines computed by the model when the position of the MCCs, U_0_, an polarity order from experiment are used to initialise the numerical model. The numerical model reproduces the obbserved streamlines with a good agreement. Comparison is shown for two differents heights (*z* = 30 µm and *z* = 90 µm). The maps of angular errors quantify the orientation difference (in degree) between the local flow velocity vectors measured via PIV and the flow velocity vectors computed.(C) Comparison of the average velocity flow profiles between experiments (5 explants) and simulations. Although the simulation grid spacing is 20 *µ*m, we use interpolation to obtain values every 10 *µ*m, allowing comparison with experimental measurements.

The computational domain includes a solid wall at the bottom, representing the epithelial surface, on which a no-slip boundary condition for the fluid velocity is imposed. The four lateral walls in the x and y directions are modeled as open boundaries to ensure the fluid flow is unrestricted by artificial computational barriers. The domain size was optimized using a sensitivity analysis to ensure that boundary proximity had negligible effects on the numerical results (Fig. S6). For consistency and to ensure grid-converged results, a uniform grid size of *dx* = *dy* = *dz* = 20 µm was used across all simulations.

#### Validation of the model

We validated our model by comparing the numerical flow fields with experimental measurements obtained in untreated and axitinib-treated tissue explants. Figure 4B presents a side-by-side comparison of simulated and experimental streamlines at heights 30 and 90 µm for a representative explant (n°9, see Fig. S4). The local agreement is quantified by color maps of the angular error, defined as the local difference in flow direction between the simulation and experiments. Furthermore, Figure 4C compares the average planar velocity magnitudes across multiple explants at various heights. Note that vertical velocities were excluded from comparison due to the current inability to measure them experimentally.

Overall, there is good agreement between the model and experimental data. Streamlines from both simulations and experiments visually demonstrate a consistent lateral flow across the entire explant domain. Close to the tissue, significant but highly localized angular errors are observed. These errors are likely attributable to small, unresolved tissue folds that harbor motile cilia. At greater distances from the tissue surface, these angular errors become minor, confirming that the model accurately reproduces the large-scale flow directions. Simulated average planar velocities also show very good agreement for all explants, with a deviation of less than 10% from experimental measurements (Figure 4C).

These results validate our simulation approach as an effective tool for describing fluid flows at the surface of ciliated epithelia. In the following section, we leverage this validated model to analyze flow over embryonic flanks, operating under the assumption that the tissue’s slight convexity does not significantly alter the established flow patterns.

### 4. Computing 3D flows on embryo flanks

Leveraging this validated numerical model at hand, we proceeded to predict flow patterns at the surface of *Xenopus* embryo flanks. Although the embryo is weakly curved, its curvature is negligible at the scale of the flank. Therefore we reasonably approximated the epithelial surface as planar in our simulations.

#### Initial conditions for the model

The static input parameters, such as MCC position and distribution, were directly extracted from images of intact embryos. As MCC beat directions cannot be measured directly on fixed embryos, we adopted an average level of alignment based on explant tissue data, corresponding to a polarity order of 0.6 (which falls within the observed range of 0.4 to 0.8 at a radius of 500 micrometers for explants). To establish this polarity, beat directions were initially assigned randomly to each MCC and then iteratively adjusted until two criteria were met: (i) achieving a polarity order of 0.6 at a 500 µm radius and, (ii) reproducing the in-vivo major streamlines observed on the embryonic surface (see Figures 5A & B). Finally, the characteristic velocity *U*_0_ for each MCC was randomly assigned following a Gaussian distribution with a mean of 350 µm/s and a standard deviation of 250 µm/s. These parameters were chosen based on explant measurements, where velocities ranged from 20 µm/s to 800 µm/s in a Gaussian-like distribution. As shown in the horizontal projection of the streamlines (Figure 5B), the model successfully reproduces the characteristic diagonal flow (from anterior-dorsal to posterior-ventral) and the distinct streamline patterns on the dorsal side of the embryo (4).

**Fig. 5.**
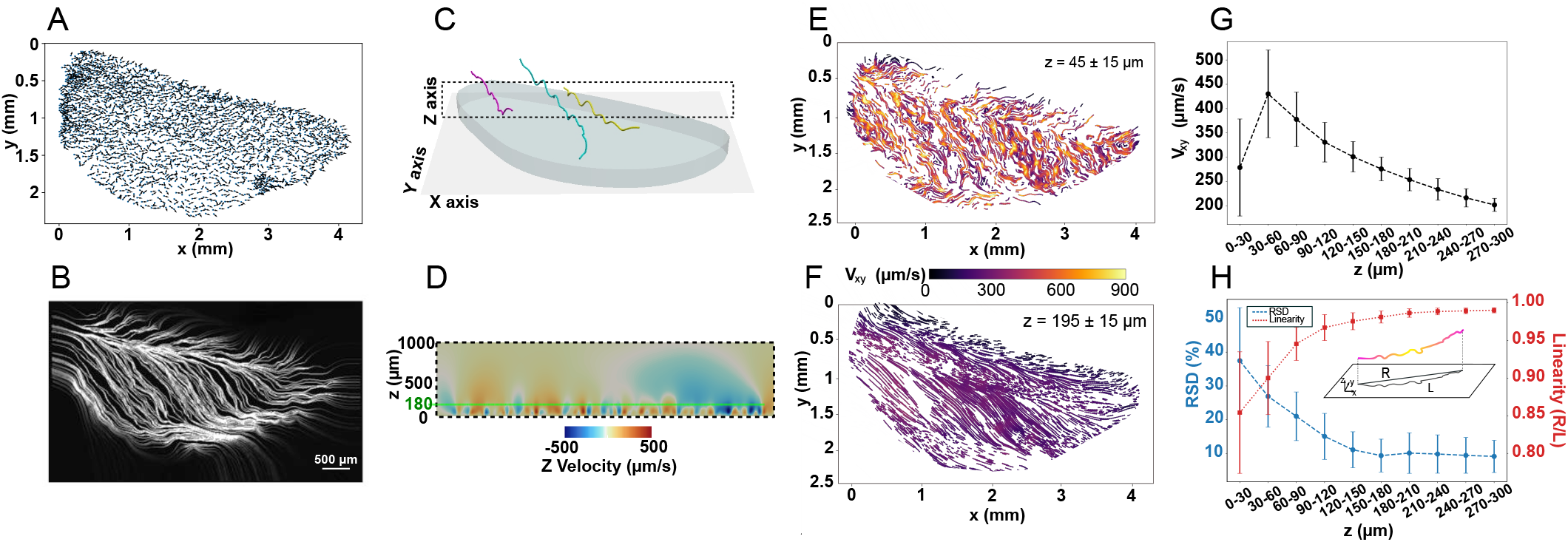
Characterisation of simulated streamlines at the embryo’s scale. (A) Plotting of MCC orientation vectors on a virtual embryo to achieve an alignment order of 0.6 at 500 µm range. (B) Numerical streamlines projected in 2D, reproducing classical features observed experimentally, with an anterior–posterior flux along the flank. (C) Example of three numerically predicted streamlines showing anterior–posterior trajectories with fluctuations along the *z*-axis. The (*x, z*) dotted line plane corresponds to the cross section depicted in panel D. (D) Cross-section of the embryonic flank in the (*x*–*z*) plane with *Z*-velocity shown as a colormap, highlighting alternating regions of upward (red) and downward (blue) flow, especially for *z <* 180 µm. (E) Projected streamlines and planar velocity as colormap for *z* = 45 µm (+ or-15 µm). (F) Projected streamlines and planar velocity as colormap for *z* = 195 µm (+ or-15 µm). Only partial streamline segments are captured due to fluctuations along *z*. (G) Mean planar velocity along streamlines as a function of height. (H) Relative standard deviation (RSD) of planar velocity and linearity as a function of height. The insert shows a streamline, with *R* and *L* definition. Linearity is defined as the ratio *R/L*. The color changes along the streamline represents the planar velocity variations, which are directly related to the RSD value. Error bars represent the standard deviation across the streamline population.

#### Flow characterization and spatial analysis

The resulting flow was characterized by analyzing the individual streamlines.

On the embryo flank, the streamlines are predominantly oriented along the horizontal anterior–posterior axis (Fig. 5C). Variations in vertical height (along the *z*-axis) are primarily due to recirculation patterns in the plane. This is illustrated in Figure 5D, where the z-velocity is mapped, showing upward (red) and downward (blue) flow. The strongest z-velocities are observed close to the epithelial surface, at heights below 180 µm. However, importantly, the mean *z*-velocity along the streamlines is close to zero (Fig. S7), supporting the conclusion that the bulk flow exhibits an average two-dimensional character.

Figures 5E & F show the projected streamlines at two different heights, z = 45 µm (+ or-15 µm) and *z* = 195 µm (+ or-15 µm). The color scale indicates the velocity magnitude in the epithelial plane. Since streamlines fluctuate in *z*, only short segments (typically spanning about 1 mm) are captured at a given height. Consistent with explant observations, streamlines become less tortuous and flow velocities decrease with distance from the epithelial surface. Concurrently, the variability in velocity along a streamline also diminishes with height. Quantitative results (Figures 5G & H) display the variation in streamline linearity, defined as the ratio of end-to-end displacement (R) to total length (L) (see insert of Figure 5H), mean planar velocity, and its relative standard deviation (RSD), an indicator of the velocity heterogeneity, as a function of height. The plateau observed in both the RSD of planar velocity and the streamline linearity at a distance of approximately 180 µm allows us to define two distinct flow regions: a Near-Field region (below 180 µm), where recirculation induces tortuous streamlines and inhomogeneous planar velocity, and a Far-Field region (above 180 µm), where the flow becomes more planar and uniform.

#### Parametric analysis: effects of simulation input parameters on flow characteristics

To understand how biological variability influences the flow driven by MCCs on the embryonic flanks, we investigated the impact of four key conditions: the characteristic velocity (*U*_0_) of MCCs, the MCC density, their spatial organization (order parameter) and their beat direction alignment (polarity order at a 500 µm radius). The baseline conditions were *U*_0_ = 350 *µ*m/s, a density of 3.2 ·10^−4^ MCC/µm^2^, order parameter of 0.6, and polarity order of 0.6.

Each condition was varied as follows:

- The characteristic velocity (*U*_0_) was reduced to 200 µm/s and 100 µm/s.
- The MCC density was varied to include a low-density scenario (1.6 *·* 10^−4^ MCC/µm^2^) and a high-density scenario (6.5 *·* 10^−4^ MCC/µm^2^).
- The spatial organization was varied by introducing MCC clustering, lowering the order parameter to 0.45, or creating an MCC-free zone (size 1 mm^2^) near the head region, which also resulted in a reduced order parameter of 0.2.
- The beat direction alignment was varied to include a low-polarity scenario (polarity order of 0.4) and a high-polarity scenario (polarity order of 0.8).

Figure 6 presents the mean planar velocity across all tested conditions at *z* =45 µm (dark band, representing the tortuous, near-field region) and *z* = 195 µm (light band, representing the uniform, far-field flow). The variations in RSD and linearity are presented in Supplementary Fig. S8. Qualitatively, the observed behaviors are similar at both tested *z*-values. The linearity of the streamlines is primarily influenced by the polarity order, while being relatively unaffected by other conditions (see Fig. S8). Increasing *U*_0_ yields a significant increase in planar velocities (Fig.6A), without altering the RSD (Fig. S8). Increasing MCC density leads to a modest rise in planar velocities (Fig. 6B) and contributes to greater velocity homogeneity (reduced RSD, Fig. S8). However, above a critical density, the velocity saturates. This behavior is expected, when the average velocity close to the epithelium converge to the intrinsic velocity of MCCs *U*_0_, adding more MCC doesn’t change the global flow. Increasing the polarity order raises planar velocities (Fig. 6D) and improves velocity homogeneity. In contrast, variations in the geometrical order parameter have no significant effect on the average streamline metrics (Fig. 6C). These findings suggest that flow velocity is primarily governed by the total ciliary force, density and beat direction alignment. Strikingly, the system appears robust to moderate levels of spatial disorder or small tissue defects.

**Fig. 6.**
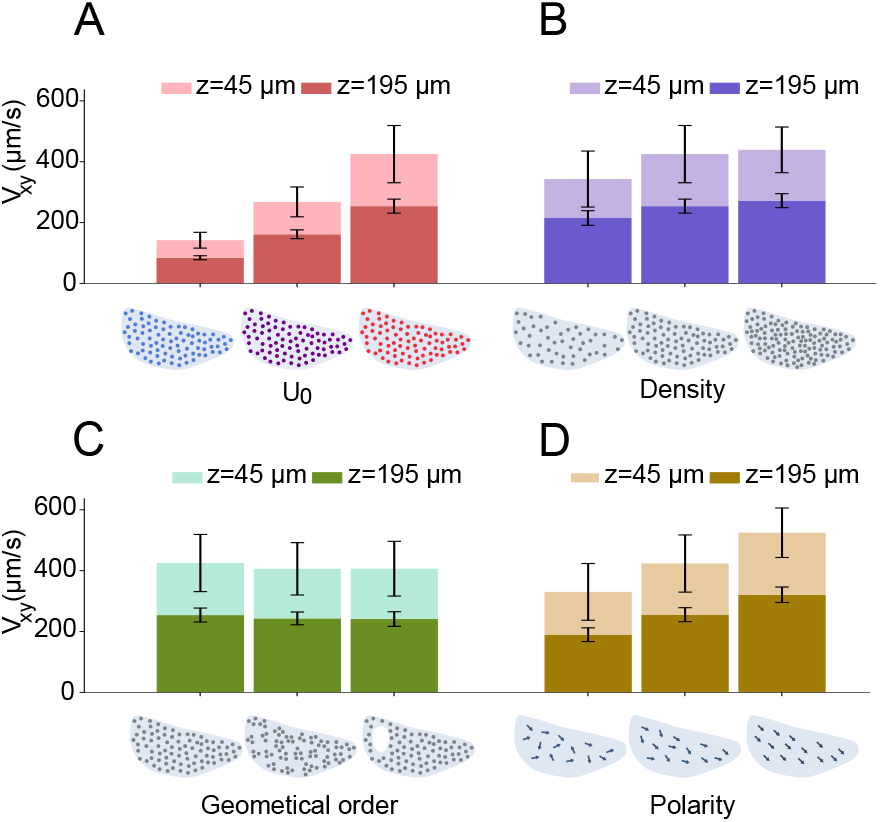
The mean planar velocity was computed for two representative z-slices: nearfield region (z = 45 µm) and far-field region (z = 195 µm), under various conditions illustrated schematically under the graphs.(A) Characteristic velocity *U*_0_ at 100, 200, and 350 µm/s. (B) MCC density at low, medium, and high levels. (C) Geometrical order for ordered, clustered, and hole configurations. (D) Polarity order for low, medium and high MCC alignment. Error bars represent the standard deviation across the streamline population

### 5. Bacterial motion and infectability

To address epithelial clearance and its robustness against infection, we computed trajectories of virtual swimming bacteria, modeled as point particles advected by the cilia-driven flow. Although bacteria can actively swim in the horizontal plane, their intrinsic swimming speed is negligible compared to the fluid velocities generated by ciliary beating. We therefore assume that bacterial motion parallel to the epithelial surface is entirely determined by the local fluid streamlines. However, active vertical motion is critical: if a bacterium swims toward the epithelium, for example via chemotaxis, this velocity must be accounted for, as the mean fluid z-velocity is zero. To account for active swimming, we introduce a tunable constant downward velocity, denoted *v*_bac_. Bacterial motion was modeled by adding a constant vertical swimming velocity to the simulated fluid velocity field, and trajectories were computed as streamlines of the resulting effective flow.

We focus our analysis on bacteria moving above the embryo’s head, as this region maximizes the transit time over the surface. This represents a worst-case scenario, offering the greatest opportunity for bacteria to reach and infect the epithelial surface. Figure 7A schematically represents the bacterial trajectories, showing the initial height (*z*_0_) and the two possible outcomes: contact with the embryo (red) or cleared by the flow (grey). Contact with the epithelial surface is defined as bacteria reaching *z* = 6 µm, corresponding to the thickness of the mucus layer. In this case, only 3 out of 35 bacteria starting at *z*_0_ = 90 µm with velocity *v*_*bac*_ = 5 µm/s reached the epithelial surface (Fig. 7A). These data were obtained from flow simulations conducted with baseline MCC conditions, *U*_0_ = 350 µm/s, a density of 3.2 · 10^−4^ MCC/µm^2^, an order parameter of 0.6, and a polarity order of 0.6. The contact probability — defined as the fraction of bacteria reaching the epithelial surface— is shown in Figure 7B as a function of their initial *z*_0_, for three distinct bacterial velocities (*v*_bac_ = 0, 2, 5 µm/s). This latter value is estimated to be among the highest for the vertical component of bacterial motility under shear flow (24–26). The probability of reaching the epithelium is high for *z*_0_ < 40 µm, which is the region where ciliary motion induces significant fluid mixing between flow layers. Above 40 µm, the contact probability decreases slowly below 10% with *z*_0_, becoming zero at *z*_0_= 150 µm, for non-motile bacteria *v*_bac_ = 0 (Figure 7B &C). In contrast, higher bacterial downward velocities consistently result in increased contact probabilities across all starting heights, maintaining a non-zero probability even at large *z*_0_. This highlights that motile bacteria can reach the epithelium, whereas non-motile bacteria are efficiently cleared by the fluid flow.

**Fig. 7.**
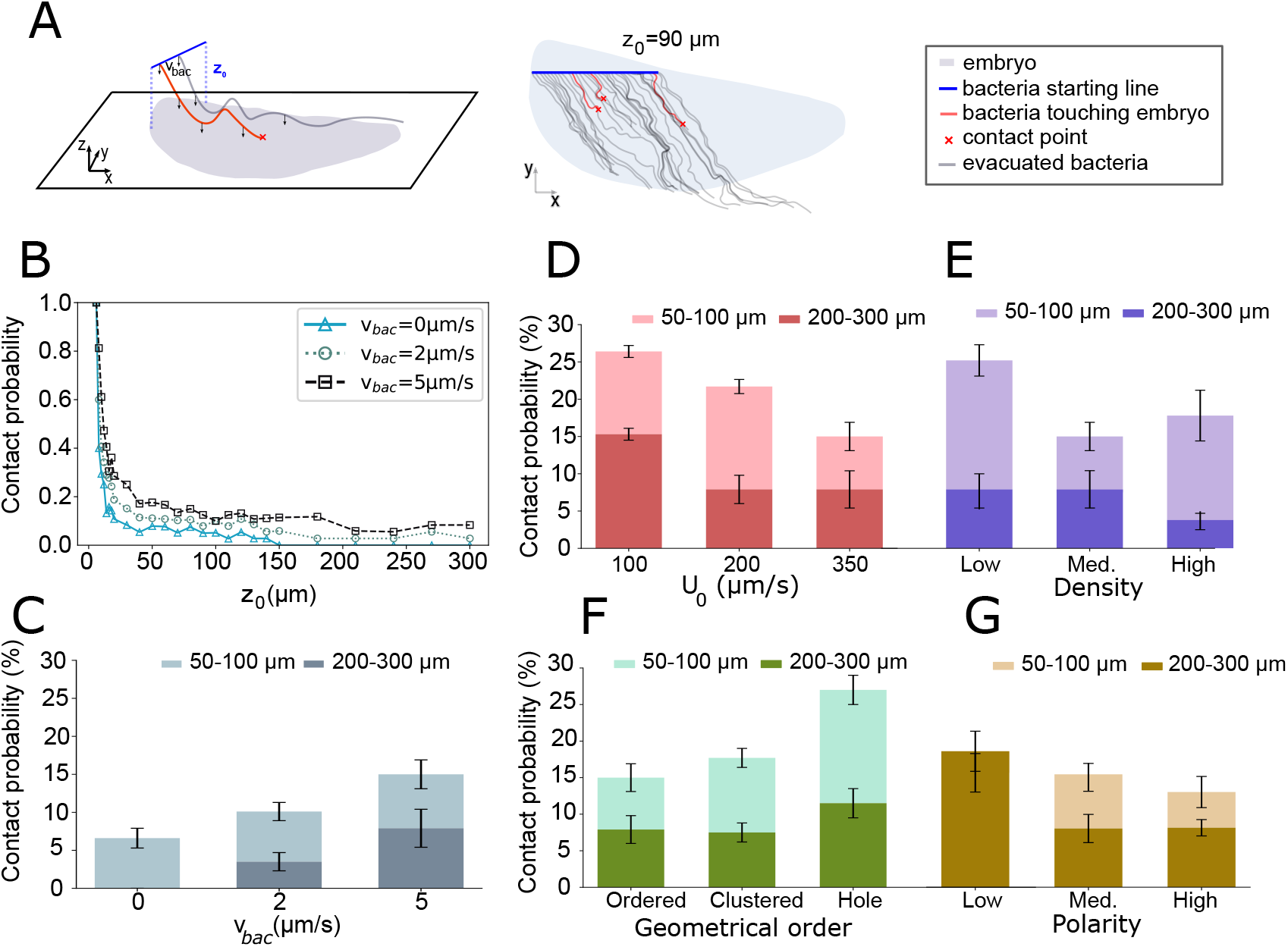
Parametric study of bacteria clearance by cilia-powered flow. (A) Schematized bacterial trajectories starting at an initial height *z*_0_ above the embryo surface (left). A 2D projection of 35 bacterial trajectories starting at height *z*_0_ = 90 µm with a bacterial speed of *v*_*bac*_=5µm/s (right). Red trajectories indicate contact with the embryo; grey trajectories indicate successful clearance. (B&C) Contact probability as a function of the initial starting height *z*_0_ for three different bacterial speeds (*v*_*bac*_). Flow simulations were conducted with baseline MCC conditions, *U*_0_ = 350 *µ*m/s, a density of 3.2 10^*−*4^ MCC/µm^2^, an order parameter of 0.6, and a polarity order of 0.6 (B) Contact probability as a function of *z*_0_ value. (C) Mean contact probability in 50 100 µm *z*_0_ range (light band) and 200 300 µm *z*_0_ range (dark band). Error bars correspond to the standard deviation of contact probabilities computed within each *z*_0_ range. (D – G) Contact probability for various tested conditions and a bacterial velocity of *v*_*bac*_ = 5 µm/s in the two *z*_0_ ranges. (D) Characteristic velocity *U*_0_ at 100, 200, and 350 µm/s. (E) MCC density at low, medium, and high levels. (F) Geometrical order for ordered, clustered, and hole configurations. (G) Polarity order for low, medium and high MCC alignment. For low polarity order conditions, the contact probabilities for both z ranges overlap. Error bars correspond to the standard deviation of contact probabilities computed within each *z*_0_ range.

The influence of ciliary velocity, *U*_0_, MCC density, their spatial distribution and their beat direction alignment on the contact probability is illustrated in Figure 7D-G for motile bacteria with velocity *v*_*bac*_ = 5 µm/s. This analysis clearly shows that *U*_0_ is a strong protective factor, significantly reducing contact probability. Conversely, the spatial distribution (order parameter) has a limited effect; only the introduction of a large defect (hole) near the head region, which disrupts the local flow, has a noticeable impact on clearance (Fig.7F). A low density of MCC results in a significant increase of contact probability, while higher densities provide a better protection (Fig.7E). The polarity order also influences contact probability, with lower polarity (i.e. low beat direction alignment) associated with a higher contact probability (Fig.7G). Together, these results show that bacterial contact with the epithelial surface is primarily controlled by the overall strength of the cilia-driven flow. Even in the presence of active bacterial motility toward the epithelium, sufficiently strong flows markedly reduce contact probability over a wide range of initial conditions.

## Discussion

Three key points emerge from our results and warrant detailed discussion.

First, we address the seemingly contradictory observation of the high survival rate of healthy embryos despite the arrival of some environmental bacteria onto the epithelial surface. Indeed, our physical model shows that even for bacteria lacking active movement towards the epithelium, the mixing of fluid layers near the surface (< 150 *µ*m) allows a proportion of bacteria to contact the epithelial surface. We can estimate the rate of bacteria landing on the epithelium per unit of time for a typical embryo (MCC density = 3.2 · 10^−4^ MCC/*µ*m^2^, *U*_0_ = 350, *µ*m/s, polarity = 0.6, and order = 0.6). The fluid volume passing over the embryo at its head (at a height z of 300 *µ*m) per unit of time is obtained by integrating the flow rate per 30-*µ*m layer, based on numerical data. Assuming a typical bacterial concentration in pond water of 10^7^ bacteria/ml (27), we deduce the bacterial arrival rate from our simulations. For non-vertically motile bacteria, this number is on the order of 50 bacteria/s. Considering that only 10% of these might be pathogenic, approximately 5 pathogenic bacteria per second land on each flank of the embryo. Given a flank surface area of about 5 mm^2^, this equates to 1 bacteria · mm^−2^ s^−1^. These arriving bacteria first contact the 6-*µ*m-thick mucus layer covering the embryo’s skin, which possesses known antibacterial properties (17). Their subsequent elimination upon arrival provides a first, albeit coarse, estimate of the bacterial load that the mucus is capable of clearing per unit of time. The strong ciliary flow is coupled with the action of the mucus layer to successfully clean the surface and eliminate the arriving microbial load.

The second addressed point concerns the initial question of whether the physical system is optimized for maximum clearance. Our results show that substantial perturbations to the spatial distribution of MCCs do not significantly affect bacterial clearance. Therefore, this specific cellular arrangement is not optimized to maximize the clearance rate. More generally, the system’s characteristics (*U*_0_, MCC density, polarity, and order) must be perturbed quite significantly to substantially impact the probability of bacterial contact. The biological system does not seek to optimize these parameters for maximum clearance but rather to ensure high robustness. This robustness allows embryos to develop without infection even if their physiological characteristics are somewhat degraded or if the environment is more hostile (e.g., higher bacterial concentration). This observation suggests that optimizing ‘performance’ metrics—such as speed or energy efficiency—may not be the primary guiding principle for the biological systems investigated here. Instead, these systems prioritize the ability to maintain function and survive even under compromised conditions, such as those resulting from injury, environmental stress, or developmental issues. Finally, we reflect on the functional implications of the highly homogeneous spatial distribution of multiciliated cells, which is conserved feature in the embryonic skin of amphibian species (2). Our data suggest that the global flow powered by MCC collectives, and the associated clearance capacity, are not dependent on MCC geometrical order. It remains possible, however, that regular MCC distribution matters at a more local scale, and closer to the skin. Indeed, our data reveal fluctuating non-zero vertical flow velocities near ciliary bundles, that may optimize oxygen uptake. For such function, regular MCC spacing would appear as an optimal solution. Moreover, MCC distribution in the superficial cell layer is expected to indirectly impact the distribution of secretory cells, which may also be an important factor for functional mucociliary protection. A final potential implication that we would like to mention is linked to the death of MCCs at pre-metamorphic stages. Death and removal of cells in epithelia represent a challenge, as junctions with surrounding cells are temporarily lost and must be rebuilt to preserve tissue integrity (28). The concomitant death of clustered MCCs would represent a major risk of epithelium disintegration, which would be avoided through regular paving of MCCs.

In conclusion, by combining experimental measurements, on both embryos and in-vitro explants, with a computational fluid dynamics model, we contributed to a better understanding of the critical biophysical mechanisms involved in the protection of *Xenopus* embryo at early stages. Our parametric analysis revealed that the efficiency of pathogen clearance is mainly governed by the intrinsic flow velocity U_0_ of MCCs. Significantly, the system exhibits remarkable functional robustness, as clearance efficiency remains high and largely insensitive to moderate variations in MCC density, spatial arrangement, and beat direction alignment. This finding suggests that the biological mechanism prioritises system resilience and reliable protection under non-ideal or compromised conditions over the optimisation of energy consumption. The question of the role of the specific tissue patterning of MCCs remains open.

## Materials and Methods

### Xenopus laevis

All experiments were performed following the Directive 2010/63/EU of the European parliament and of the council of 22 September, 2010 on the protection of animals used for scientific purposes and approved by the “Direction départementale de la Protection des Populations, Pôle Alimentation, Santé Animale, Environnement, des Bouches du Rhône” (agreement number G 13055 21). Adult *Xenopus laevis* individuals were obtained from NASCO. Female frogs were injected with human chorionic gonadotropin a day before egg collection, and eggs were externally fertilized in a petri dish with male sperm. Embryos were then dejellied with a 3% cysteine hydrochloride solution (pH 8), rinsed, and kept in 0.1X MBS solution (Modified Barth’s Saline) at 13 °C until they reached the desired developmental stage.

### Foxj1 CRISPR/Cas9 injection

Following in vitro fertilization at 18 °C, eggs were dejellied with 2% cysteine hydrochloride solution (pH 8), rinsed a few times with tap water and once with 0.1X MBS. Eggs were then transferred to a 4% Ficoll solution at the 1-cell stage and injected with 5 nl of recombinant Cas9 (0.2 ng/nl, PNA Bio, #CP03) or Cas9 + FoxJ1 single guide RNA (0.06 ng/nl; 5’-GCAGCTAATACGACTCACTATAGGGATACATACCTGCCAGG TGTTTTAGAGCTAGAAATAGCAAG-3’)(29) near the centre of the animal pole. Injected embryos were kept at 23 °C for at least 3 hours before transferring them to 2% Ficoll at the desired temperature overnight. The next day, they were transferred back to 0.1X MBS for subsequent steps.

### Flow measurement on embryos

Tadpoles were kept in anaesthetic (0.02% MS-222 in 0.1X MBS) lying on their sides. 10 µm fluorescent beads (Invitrogen) were then added near the top of the head using a 2 µl syringe (Hamilton) and videos of the moving beads were recorded using an epifluorescence microscope (Nikon SMZ18) at 3X magnification attached with a digital camera (Hamamatsu ORCA-fusion C14440) using 10 ms exposure. A small region of interest near the middle of the tadpoles was analysed using the Track-Mate plugin (Kalman tracker) in ImageJ to quantify the mean speed of each bead.

### Imaging multiciliated cells in embryos

To reveal MCCs, embryos were fixed at stage 30 in PFA 4% in PBS-Triton 0.1% during 30 min at 23°C, rinsed 4 times in 1X PBS and blocked in BSA 3% in PBS for 1h at 23°C. Incubation with mouse anti-acetylated-Tubulin (Sigma T7451, 1/1000) was done overnight at 4°C. The next day, embryos were rinsed 4 times with 1X PBS and washed 3 times for 30 min in 1X PBS. Incubation with goat anti-mouse IgG2b-568 (Life Technologies, 1:800) was done 1h at 4°C. Embryos were rinsed 4 times in 1X PBS and washed 3 times in 1X PBS for 5min. Embryos were imaged using an epifluorescence microscope (Nikon SMZ18) at 3X magnification attached with a digital camera (Hamamatsu ORCA-fusion C14440).

For quantitative analyses of MCC spatial organization, eight control embryos and twelve axitinib-treated embryos (see Axitinib treatment below) were fixed and imaged at NF stages 29–30. For each embryo, the flank epidermis was imaged over surface areas ranging from 3.18 to 5.10 mm^2^. These datasets were used to extract MCC positions for subsequent quantification of spatial organization and geometrical order.

### Quantification of infection rates by *Aeromonas hydrophila*

*A. hydrophila* (Chester Stanier, ATCC®7966; recombinant expressing GFP and kanamycin resistance) was kindly provided by Eamon Dubaissi (University of Manchester, UK). Stock colonies were maintained on LB agar plates supplemented with 50 µg/ml kanamycin at 4 °C for a month. *A. hydrophila* was freshly grown in Tryptic Soy Broth (TSB) liquid culture (50ml) supplemented with kanamycin (50 µg/ml) to OD600=0.8 to 1 (30°C, 232rpm). Bacteria were pelleted by centrifugation 10min at 3500xg, washed in MBS 0.1x to remove traces of TSB, pelleted again and resuspended in MBS 0.1x supplemented with Kanamycin 50 µg/ml to OD=0.8 (approximately 5.10^8^ bacteria/mL). Stage 30 embryos were exposed to 3ml of bacteria suspension (5 embryos/well in 12-well dishes) for 9h at 23°C. At the end of the infection period, embryos were extensively washed to eliminate bacteria in suspension (5 times in 50ml tap water), and individually crushed in 500ml TSB in 1.5ml Eppendorf tubes with a tissue homogenizer. 100ml of embryo lysate was spread on a LB Agar plate supplemented with kanamycin 50µg/ml. Plates were incubated overnight at 37°C, after which colonies were manually counted to quantify colony-forming units per infected embryo. Individual embryos were arbitrarily considered infected at cfu>100 per embryo. Controls included plating of the initial bacterial suspension and of the culture medium at the end of the infection period to check bacterial health, and plating of the last embryo wash to verify washing efficiency.

### Explant culture

The explantation protocol was adapted from established methods (30), to ensure that the tissue remained thin, flat, and transparent. Embryos kept at 13 °C were transferred to a Petri dish containing explant culture medium (Danilchik’s for Amy (DFA): 53 mM NaCl2, 5 mM Na2CO3, 4.5 mM potassium gluconate, 32 mM sodium gluconate, 1mM CaCl2, 1mM MgSO4 buffered to pH 8.3 using 1M Bicine). At stage 16, vitelline membranes were carefully removed with tweezers, the neural cord was cut using an eyebrow knife, allowing to explant a piece of ectoderm, from which the adjoined mesodermal layer was peeled away. Explants were placed in imaging chambers made with collagen-coated coverslips and PDMS walls, bonded by plasma treatment. The chambers were pre-treated with fibronectin (20 mg/ml in 1X PBS) overnight at 4°C, blocked with 1X PBS + 1% BSA for one hour, and filled with explant culture medium containing antimycotic (100 µg/ml) to prevent contaminations. To ensure flattening of the tissue, sterilized 10 × 10 mm glass slides coated with silicone grease were gently placed on top of the explant for 12 h, then removed to avoid disrupting cilia growth. Sibling embryos were grown in parallel to monitor development and to stage explants. The culture medium was replaced daily, and all solutions and dissection tools were sterilized before use.

Explants were cultured for 3–6 days prior to imaging. We observed that explants imaged after 6 days of culture (corresponding to embryonic stages 30–32) consistently exhibited stronger and more coherent cilia-driven flows than those imaged after 5 days (Fig. S2). All quantitative flow analyses presented in this study were therefore performed on day-6 explants to ensure robust ciliary activity (see Movie S1). For these analyses, a total of nine control explants and nine axitinib-treated explants were prepared and imaged.

### Axitinib treatment

We disrupted MCC distribution in *Xenopus* embryos and explants using axitinib (Sigma), a Kit receptor chemical inhibitor known to block the cell migration responsible for their regular spacing (22). Embryos at NF stage 12 were incubated in 0.1x MBS containing 50 µM axitinib diluted in DMSO, and allowed to develop until NF stage 29–30. For explants, embryos were treated with axitinib at NF stage 9/10, then explanted at NF stage 16 and transferred to explant culture medium containing 50 µM axitinib.

### Imaging explants and Flow measurements

Observations were made using a Nikon Eclipse Ti inverted optical microscope equipped with an Andor Neo 5.5 sCMOS camera, offering a frame rate above 100 fps, sufficient to visualize ciliary beating. Images were captured using Micromanager 1.4 software with a 20X objective (CFI Plan Fluor LWD 20X;N.A. 0.70). To image flow fields, fluorescent beads (FluoSpheres Polystyrene Microspheres, 1.0 µm, red fluorescent (580/605), #F13083 thermofisher) were added to the medium at a 1:1000 dilution.

### Image analysis

For every field of view, two videos were acquired: one in bright field to visualize the dynamics of MCC and the other in fluorescence, in presence of beads, for recording the flow fields associated with those MCCs. All the image processing was done using ImageJ, JPIV and custom-made Python scripts.

Depending on explant geometry, flow measurements were performed either on large continuous flat epithelial regions or on smaller flat subregions. Large explants provided continuous flat areas of up to 3.7 mm^2^, while smaller explants were analyzed using one to three imaging tiles of 0.46 mm^2^ each. Among axitinib-treated explants, three displayed flat epithelial areas larger than 1.75 mm^2^, whereas six exhibited smaller flat regions that were analyzed over one or two tiles of 0.46 mm^2^ each.

### Quantification of MCC position and beat direction

MCC positions are identified from raw videos by computing the temporal standard deviation projection of the image stack, which highlights cilia activity as regions of maximum pixel intensity. These projections can be binarized in ImageJ, showing MCCs as white patches, with MCC position calculated as the centroid of these patches. Cilia-driven flow is confirmed to align with the underlying cilia’s beating direction (31), allowing the average beating direction of each MCC to be obtained by correlating cilia activity videos with those of bead movement. The resulting beating directions are spatially averaged across all cilia in an MCC.

### Quantification of MCC area and coverage fraction

The dynamic area of MCC is measured by segmenting regions of ciliary activity, identified as white patches in the standard deviation of video frames. The coverage fraction is calculated as the area with ciliary activity divided by the total tissue area analyzed. To compare coverage fraction of MCCs on explants (live microscopy) whith embryos (immunostaining), the dynamic area can be adjusted to reflect the apical membrane area of MCCs by applying a correction term based on cilia length, following the method outlined in (32).

### Quantification of cilia beating frequency

We compute the CBF using an in-house routine developed in python. We perform image processing on movies acquired at 150 fps. For each pixel belonging to a ciliated cell, we extract the time evolution of the intensity of the pixel. Then, we compute the power spectrum of this signal via the fast fourier transform. The CBF is the frequency with the highest energy in the power spectrum. The frequencies across pixels within a MCC are then averaged to obtain the mean ciliary beat frequency at the cell level.

### Quantification of geometrical order

To quantify the randomness in MCC distribution, a geometrical order parameter is defined based on the pairwise distances between MCCs, calculated using Delaunay triangulation (See Fig. 2 (B)). From the distribution of these distances, the mean and standard deviation are computed. The order parameter is then given by:

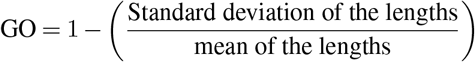

A value near zero suggests a random, spread-out distribution, while a value near one indicates a more uniform, regular arrangement of MCCs.

### Quantification of polarity order

Polarity order quantifies the alignment of beat directions of MCCs. It is determined by calculating the length of the mean resultant vector of the unit orientation vectors *p*_*i*_ for each MCC. Using circular statistics, the average of these orientation vectors provides the polarity order, with higher values indicating a more aligned beating directions. The mean 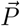 of a set of N angles (*γ*_1_, *γ*_2_, …*γ*_*N*_) is given by

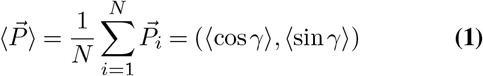

The order is then calculated as the length of the mean resultant vector

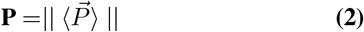

(33, 34) The order parameter reaches one when all beating directions are perfectly parallel and approaches zero for randomly oriented vectors. To measure the correlation of polarity over distance, we calculate polarity at various length scales using circles of different radii. Starting with a specific radius, polarity is computed within each circle. The radius is then incremented in discrete steps, calculating polarity for each new radius. This approach provides an average polarity value at each distance, derived from multiple circles of the same radius.

### Analysis of the flow

We quantified the flow fields via Particle Image Velocimetry (PIV), using the JPIV software. Briefly, each video frame is divided into interrogation windows, and displacement in each window is calculated as the peak of the autocorrelation function. For a 2048×2048 pixel field of view, we used 128-pixel interrogation windows with two iterations, providing a velocity vector for every 16 pixels. These fields were temporally averaged over 100 frames, equivalent to about 1 second of acquisition.

### Numerical model

We developed an in-silico model of epithelial tissue using a fluid solver, employing the threedimensional Lattice Boltzmann method for the fluid domain and integrating ciliated cells through Lagrangian points with the Immersed Boundary Method (IBM). This combined approach has been validated for simulating rigid and deformable bodies (35–37). In this study, ciliated cells are represented as point forces at height h comparable to cilia length. Three values of h were tested, and while variation was minimal, *h* = 20 µm consistently yielded the lowest mean squared error (MSE) when comparing with experimental velocities (see Supplementary S6). Input parameters include the positions (centroids of MCCs from experiments), directions (measured beating directions, and magnitudes of the point forces, derived from the characteristic velocities (*U*_0_) measured above each MCC.

The numerical method to calculate this point force *F* is the immersed boundary method. The force is:

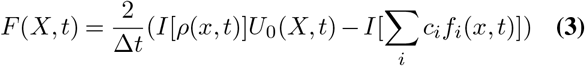

where *X* denotes the position of the MCC, *ρ* is the density of the Lattice Botlzmann fluid, *I* is an interpolating operator using numerical points surrounding the MCC to integrate values such as the density, *U*_0_ is the characteristic velocity measured above each MCC, *c*_*i*_ and *f*_*i*_ are related to the LBM (Lattice Boltzmann Method), respectively the fluid particle velocity and the fluid distribution function, *x* is the position of the LBM node and *t* is the time.

The fluid domain features a solid wall at the bottom to simulate the epithelial surface, enforcing a no-slip condition, so the fluid flow is not interrupted by any barriers. The upper wall boundary condition should mimick an inifinite media. As a compromise between accurary and computing cost, an open boundary condition has been adopted at *z* = 1 mm corresponding to 50 nodes in the vertical direction. Point forces are centered within the simulation domain to minimize boundary effects, with optimal distances between the open boundary and the epithelial domain determined through multiple trials with varying distances (Fig. S6). A grid size of *dx* = *dy* = *dz* = 20 µm was chosen, reflecting the size of the MCCs and optimized using experimental data. The fluid used in the simulation, driven by ciliated cells in *Xenopus* embryos, is water. Therefore, the density of the medium is set at *ρ* = 1000 kg/m^3^ and the kinematic viscosity (*µ*) = 1 x10^−6^m^2^/s.The Reynolds number, calculated as 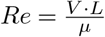, is approximately 0.01, considering a velocity of 1 mm/s over alength scale of 10 µm. Such a low *Re* indicates that the flow is in the Stokes regime. Simulations were run until a steady state was reached. All flow fields and velocity profiles shown correspond to the stationary regime.

---

**Supplementary Figure S1.**
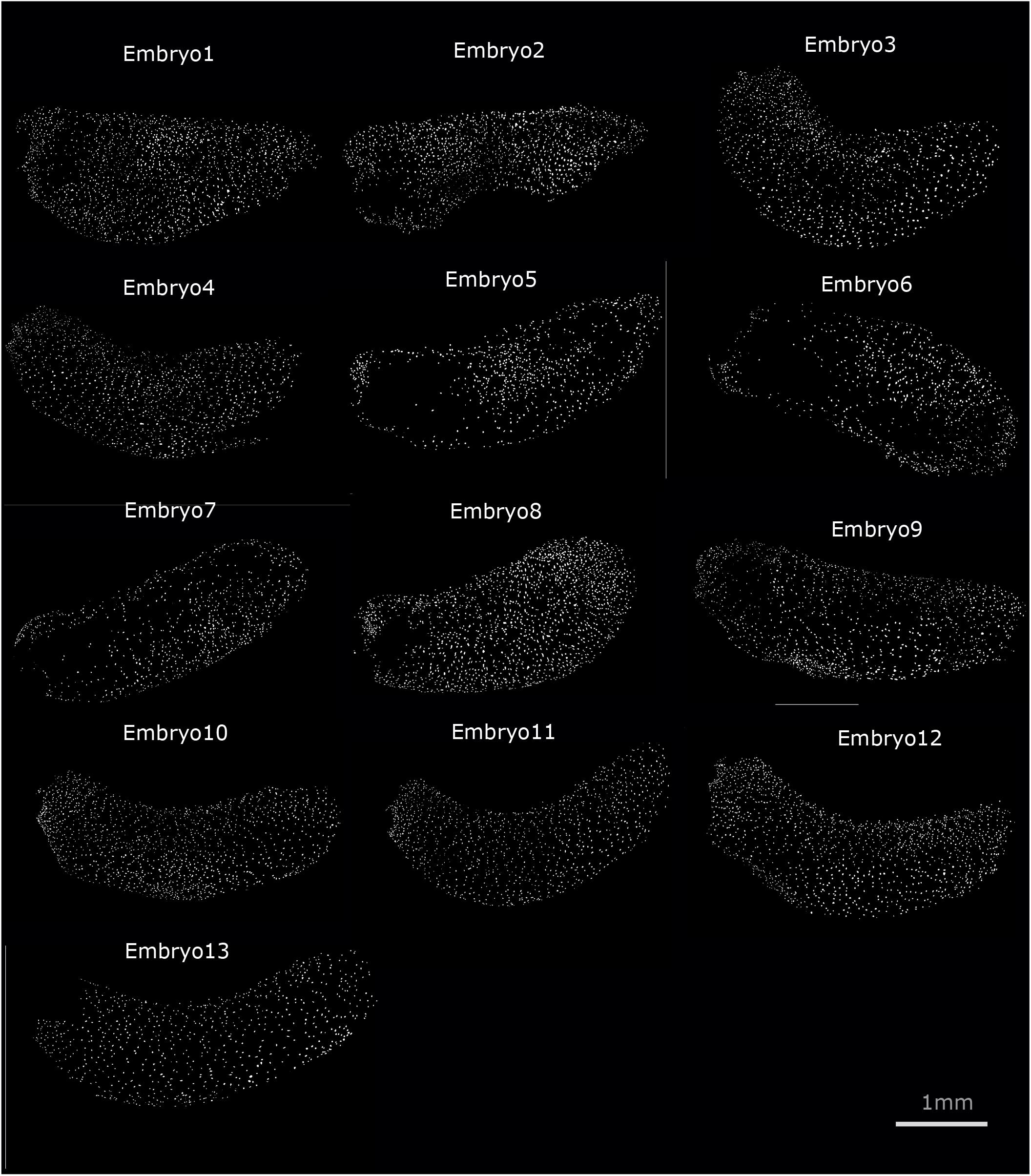
Images of Axitinib treated embryos with MCC denoted as white dots. Embryos 5-9 exhibit large regions devoided of MCC due to MCC disappearance. These are confocal images where cilia are marked by staining Ac tub and binarised to see the MCC distribution.

**Supplementary Figure S2.**
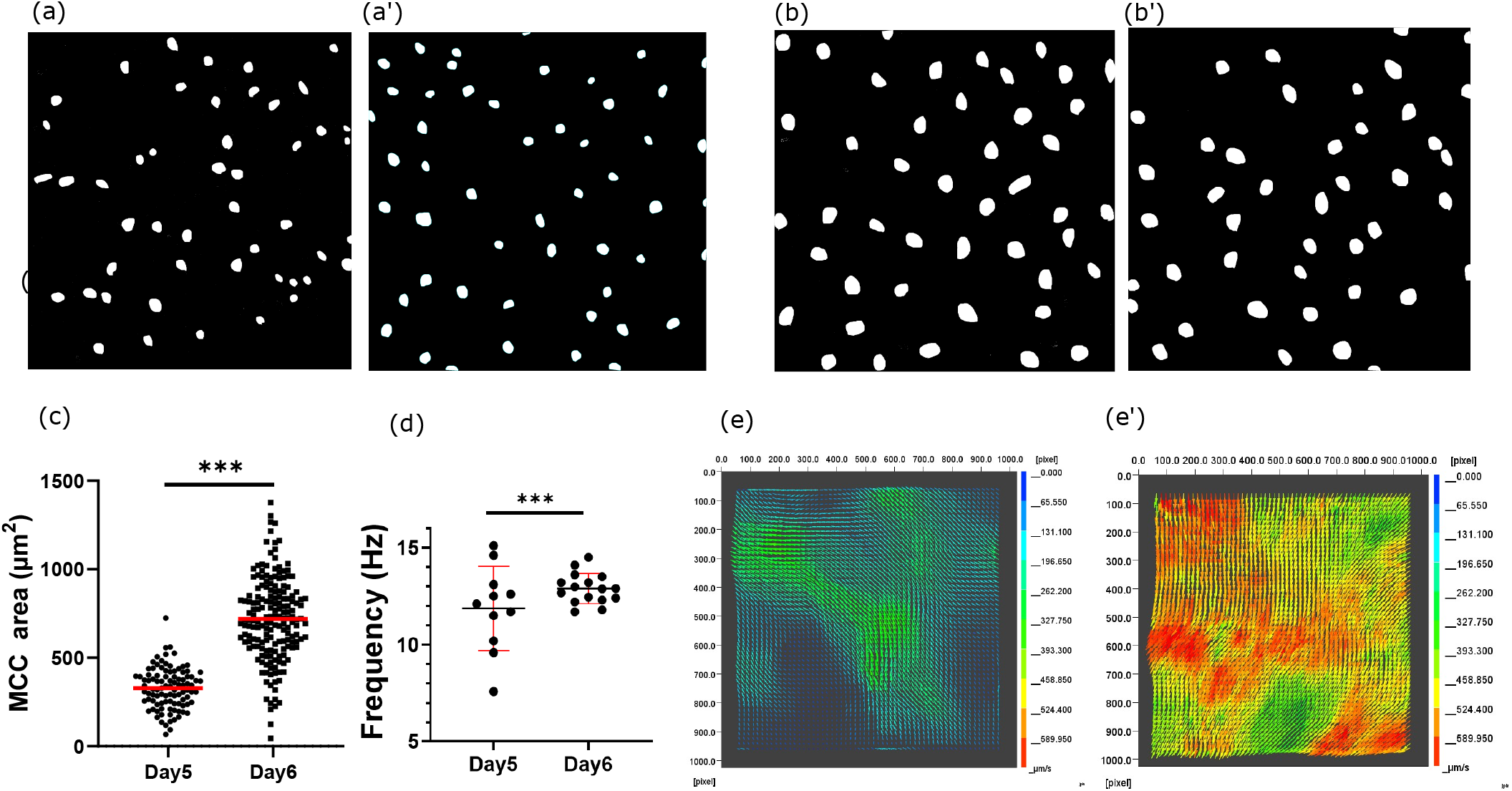
Comparison of ciliated epithelium at day 5 and day 6 of explant maturation. Fig. (a) and (a’) show ciliated cells from day 5 explants, while Fig. (b) and (b’) show ciliated cells from day 6 explants. The ciliated cell area (c) and beating frequency (d) were analyzed across multiple tiles, revealing that ciliated cells on day 6 are larger and exhibit stronger beating compared to day 5. The corresponding velocity fields generated by day 5 and day 6 explants are shown in Fig. (e) and (e’).

**Supplementary Figure S3.**
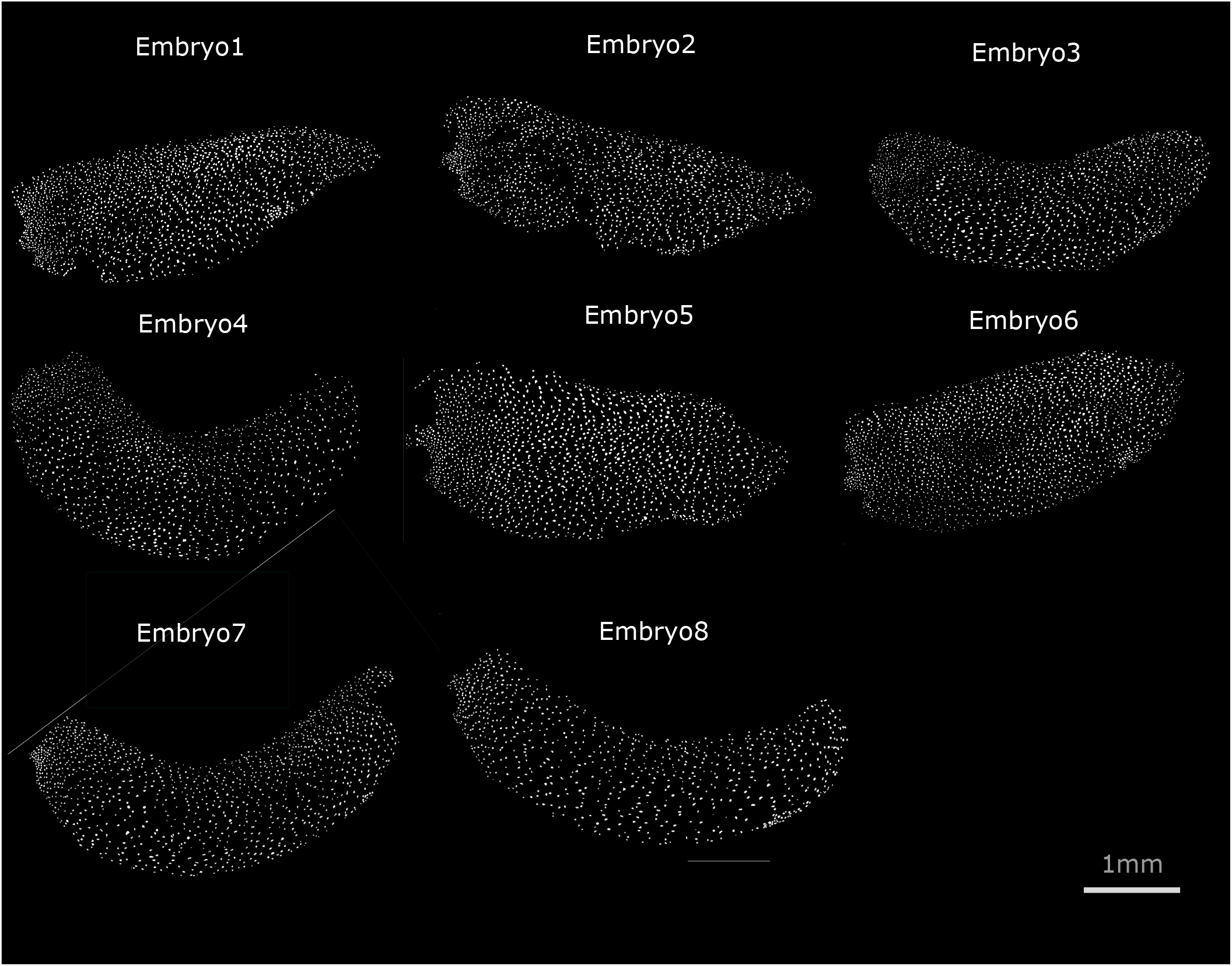
Fig. 8. Images of untreated embryos with MCC denoted as white dots. These are confocal images where cilia are marked by staining Ac tub and binarised to see the MCC distribution.

**Supplementary Figure S4.**
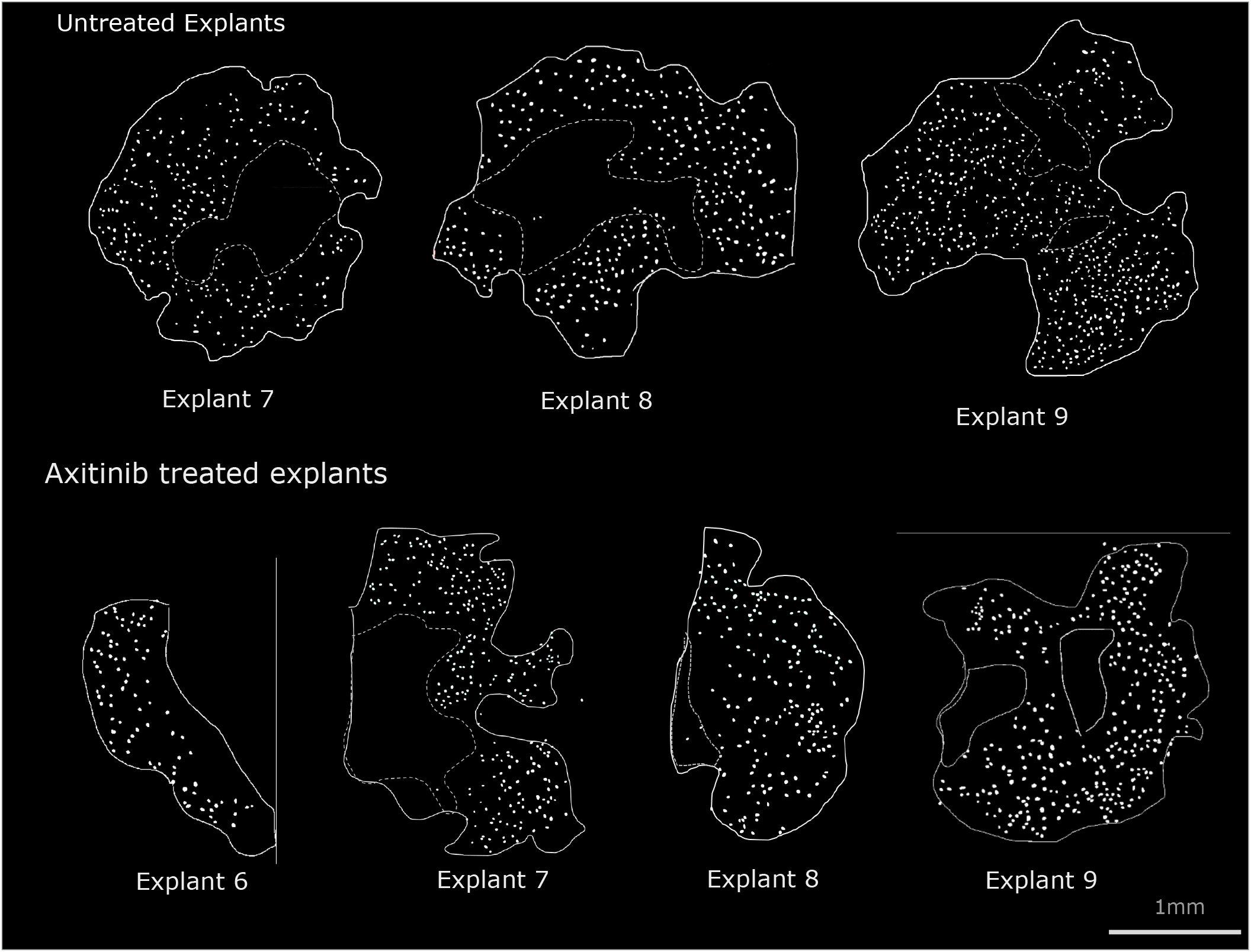
Images of explant tissues with MCC denoted as white dots

**Supplementary Figure S5.**
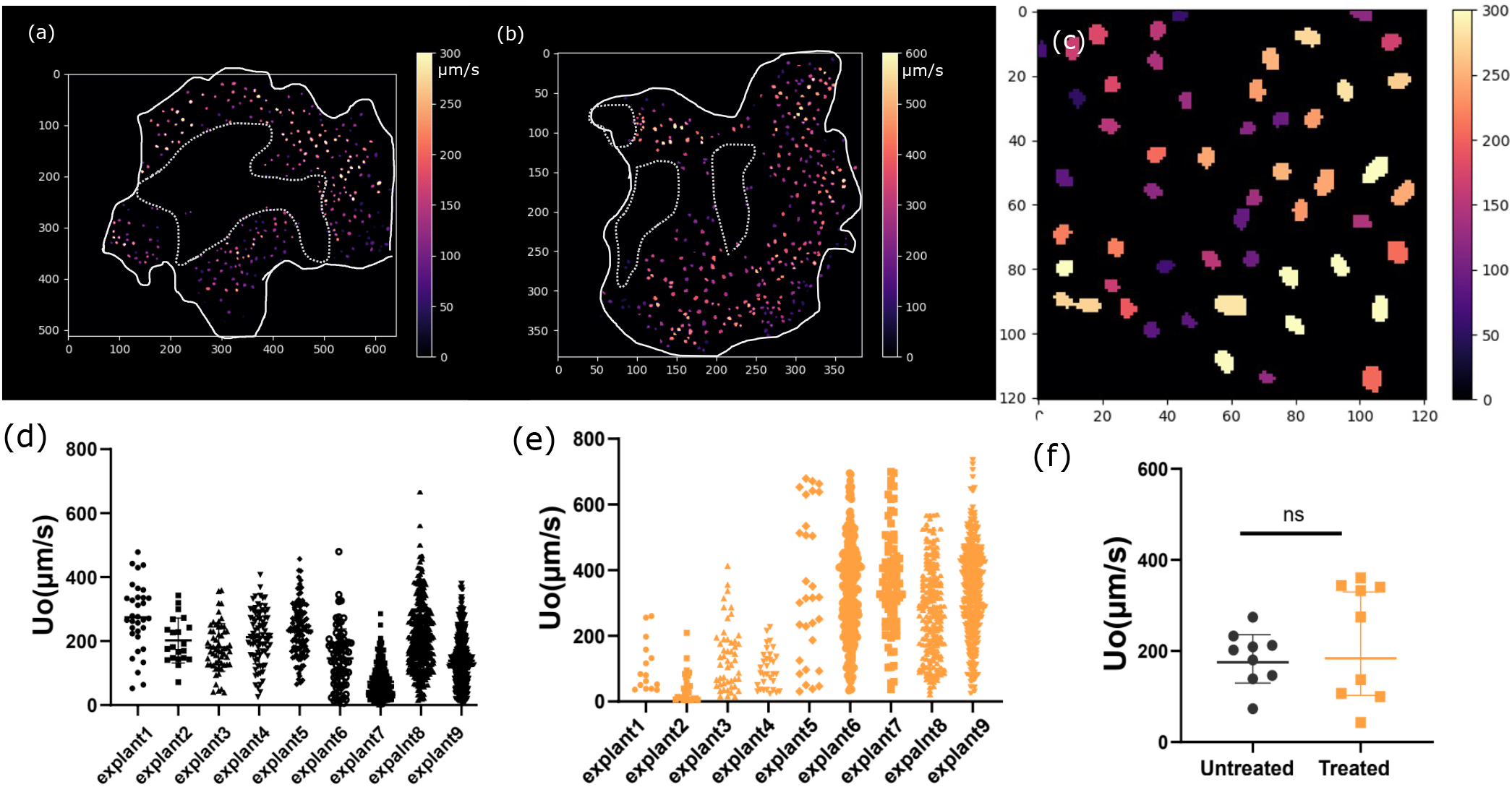
(a) *U*_0_ distribution for explant 8 under untreated conditions. (b) *U*_0_ distribution for explant 9 under treated conditions. (c) Zoomed-in image showing the variation of *U*_0_ within a tile. (d) and (e) show Uo distributions across different explants. (d) Comparison of mean Uo between untreated and Axitinib-treated explants.

**Supplementary Figure S6.**
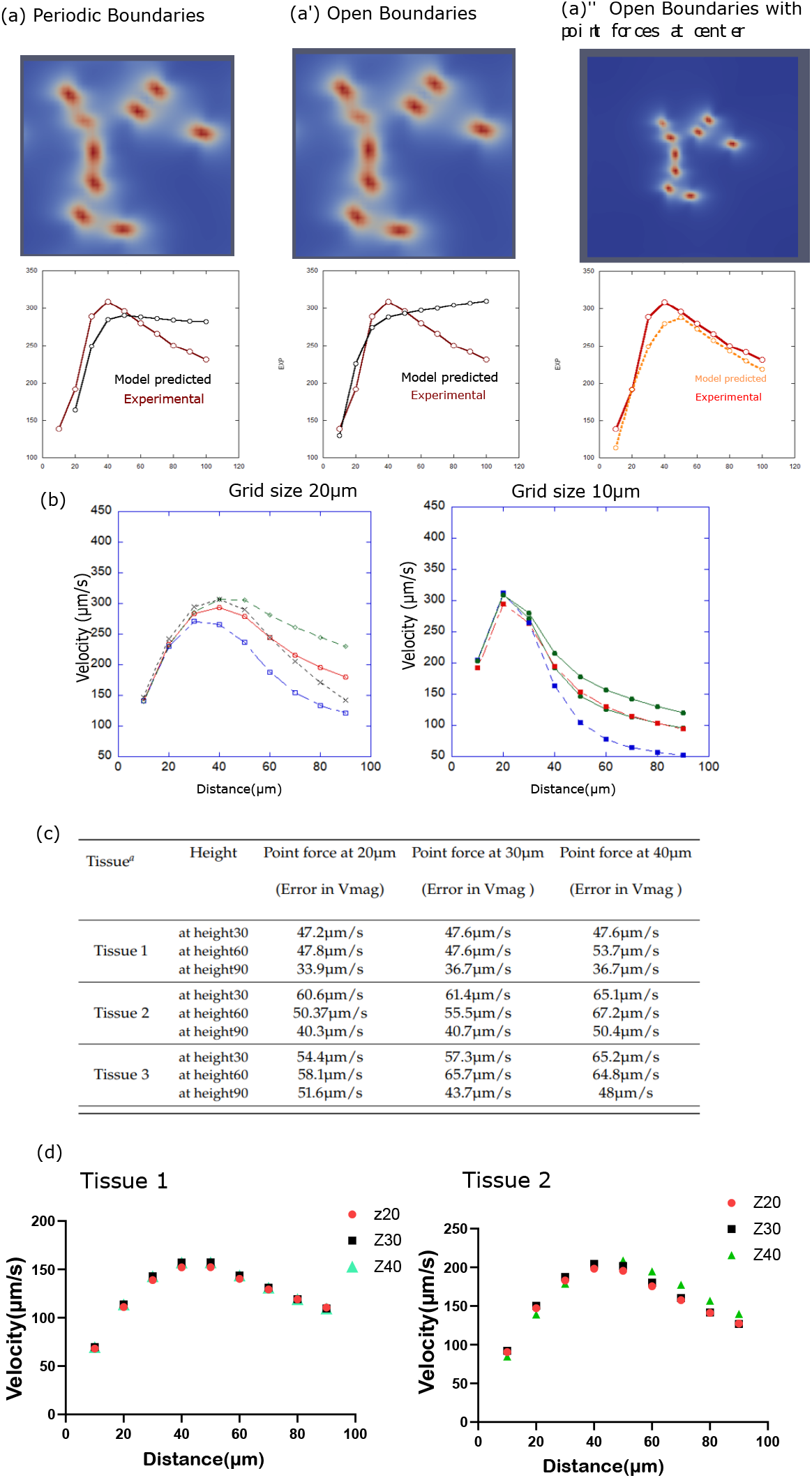
(a) Velocity profiles for different boundary conditions. The average velocity was compared across cases, with the configuration of centrally placed point forces and open boundaries showing the closest agreement with experimental results. The distance to the border was optimized through trial-and-error until boundary effects were eliminated. (b) Grid size in the Lattice Boltzmann method defined the effective size of a multiciliated cell (MCC); a grid size of 20 µm corresponded to an MCC area of 400 µm^2^, comparable to biological values. Velocity profiles above each point force were examined for grid sizes of 20 µm and 10 µm. With a 20 µm grid and point forces placed at 20 µm, the maximum velocity occurred at 40 µm, whereas at a 10 µm grid with point forces at 20 µm, the maximum velocity shifted to 20 µm due to mesh refinement. Profiles obtained with a 20 µm grid resembled experimental data more closely, and this grid size was therefore adopted. (c) Influence of the height *h*, used to represent cilia length. Three values of *h* were tested across three tissues, and while variation was minimal, *h* = 20 *µ*m consistently yielded the lowest mean squared error (MSE) compared to experimental values and was used in all subsequent analyses. (d) Velocity profiles obtained numerically for two representative tissues (left and right panels) and for the three tested values of *h*. Only minor differences between the curves were observed.

**Supplementary Figure S7.**
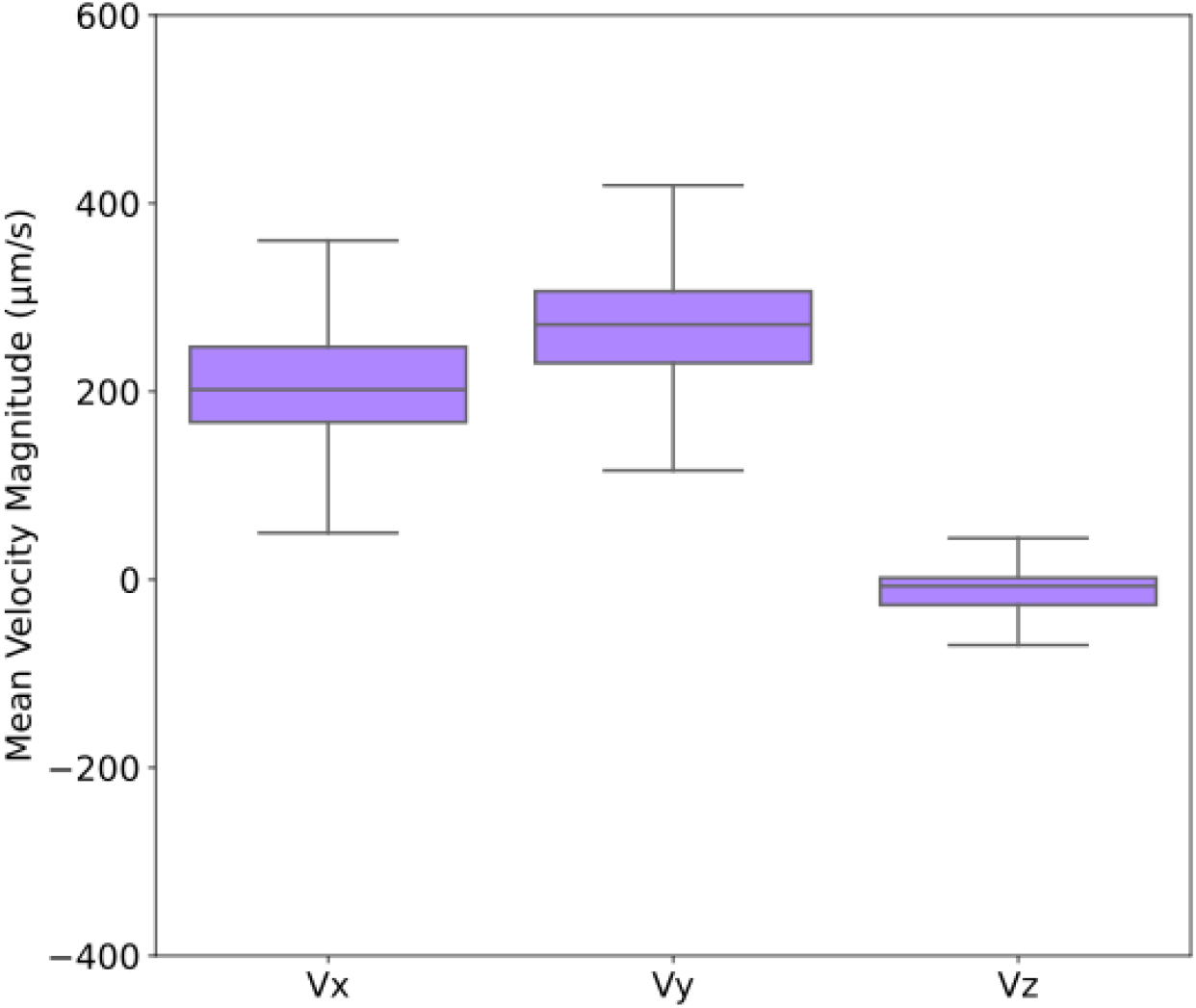
Mean componant of velocity in *x, y, z* direction on a population of approx. 400 streamlines

**Supplementary Figure S8.**
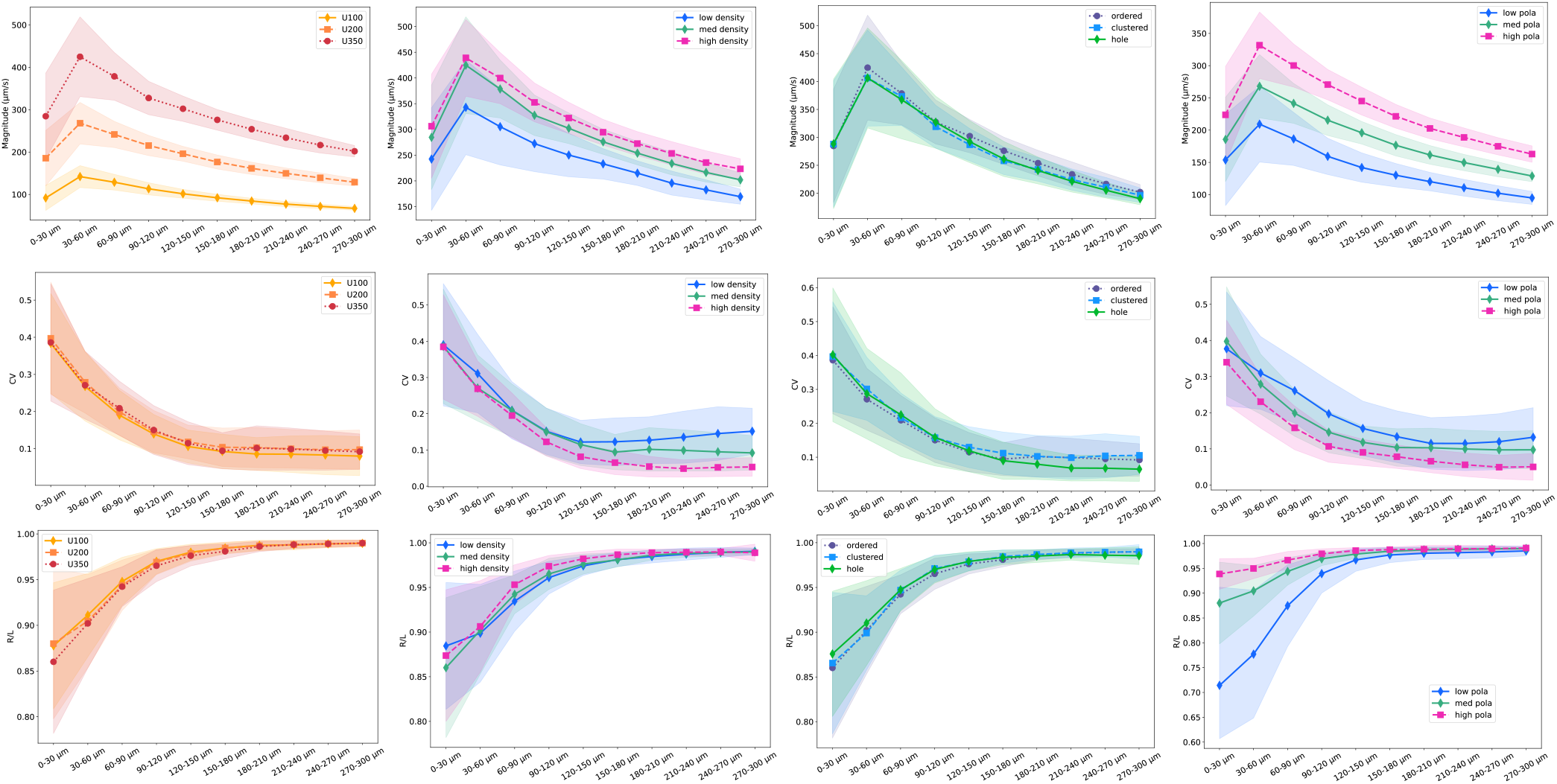
Mean planar velocity, coefficient variation (CV=RSD/100), and linearity (R/L) for all *U*_0_, density, geometrical order, and polarity conditions, plotted against the height *z* up to *z*=300 µm. Each parameter was varied starting from the baseline conditions (*U*_0_ = 350 *µ*m/s, a density of 3.2 *·* 10^−4^ MCC/µm^2^, an order parameter of 0.6 and a polarity order of 0.6), except for polarity study where the baseline conditions where (*U*_0_ = 200 *µ*m/s, a density of 3.2 *·* 10^−4^ MCC/µm^2^, an order parameter of 0.6 and a polarity order of 0.6).

**Supplementary Figure S9.**
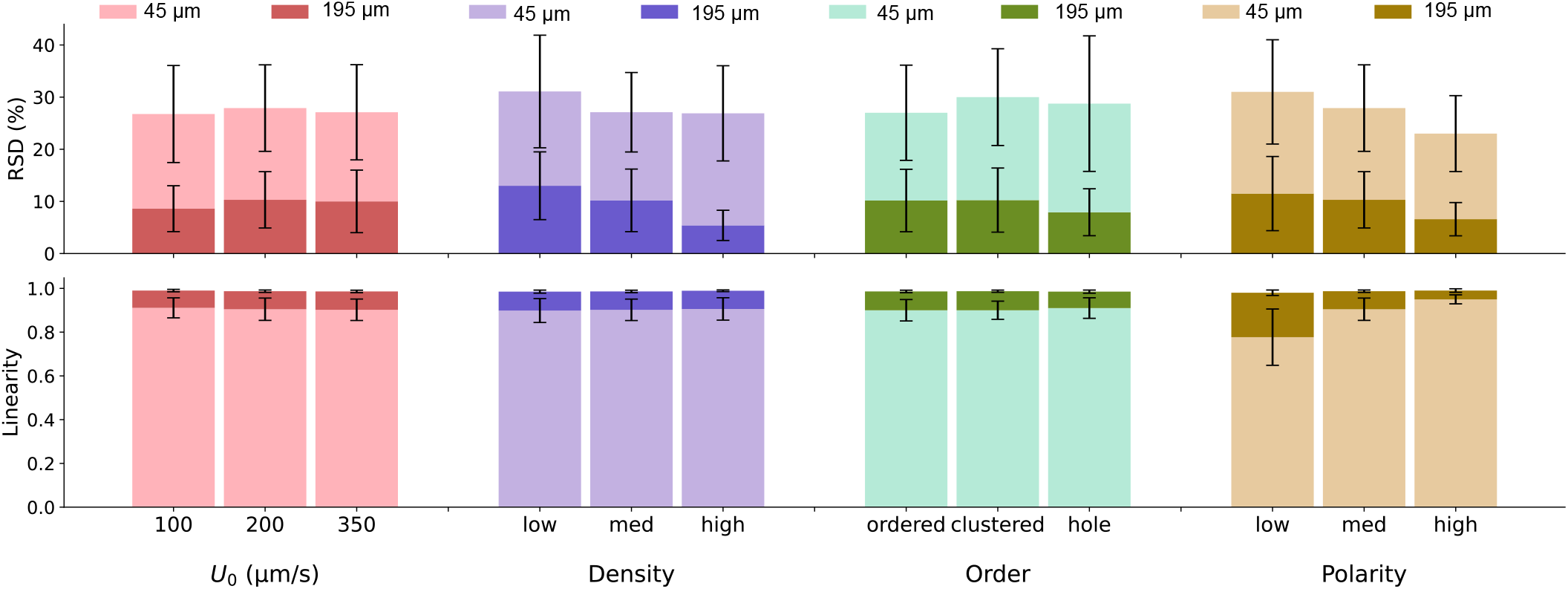
RSD and Linearity with *U*_0_, density, order and polarity variations, for two representative *z*-slices: near-field region (*z* = 45 µm) and far-field region (*z* = 195 µm). Error bars represent the standard deviation across the streamline population.

**Supplementary Table S1.**
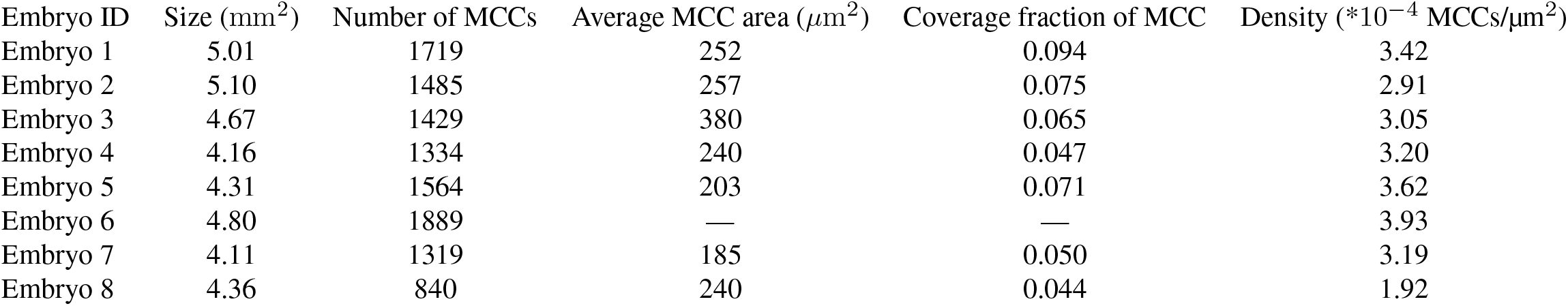
Characteristics of all embryos analyzed: Untreated. MCC area quantification for some embryos was not performed because ZO1 stained images were unavailable. Size is defined as overall area covered by the embryo.

| Embryo ID | Size (mm<sup>2</sup>) | Number of MCCs | Average MCC area (μm<sup>2</sup>) | Coverage fraction of MCC | Density (*10<sup>-4</sup> MCCs/μm<sup>2</sup>) |
| --- | --- | --- | --- | --- | --- |
| Embryo 1 | 5.01 | 1719 | 252 | 0.094 | 3.42 |
| Embryo 2 | 5.10 | 1485 | 257 | 0.075 | 2.91 |
| Embryo 3 | 4.67 | 1429 | 380 | 0.065 | 3.05 |
| Embryo 4 | 4.16 | 1334 | 240 | 0.047 | 3.20 |
| Embryo 5 | 4.31 | 1564 | 203 | 0.071 | 3.62 |
| Embryo 6 | 4.80 | 1889 | — | — | 3.93 |
| Embryo 7 | 4.11 | 1319 | 185 | 0.050 | 3.19 |
| Embryo 8 | 4.36 | 840 | 240 | 0.044 | 1.92 |

**Supplementary Table S2.** Characteristics of all embryos analyzed: Axitinib-treated. MCC area quantification for some embryos was not performed because ZO1 stained images were unavailable. Size is defined as overall area covered by the embryo.

| Embryo ID | Size(mm<sup>2</sup>) | Number of MCCs | Average MCC area (μm<sup>2</sup>) | Coverage fraction of MCC | Density (*10<sup>-4</sup> MCCs/μm<sup>2</sup>) |
| --- | --- | --- | --- | --- | --- |
| Embryo 1 | 4.85 | 1626 | — | — | 3.34 |
| Embryo 2 | 3.91 | 1338 | 222 | 0.059 | 3.42 |
| Embryo 3 | 3.97 | 1180 | 223 | 0.044 | 2.96 |
| Embryo 4 | 4.21 | 1420 | 283 | 0.040 | 3.36 |
| Embryo 5 | 5.03 | 796 | 188 | 0.035 | 1.57 |
| Embryo 6 | 4.64 | 871 | — | — | 1.86 |
| Embryo 7 | 4.35 | 854 | 170 | 0.028 | 1.96 |
| Embryo 8 | 3.97 | 1635 | 160 | 0.071 | 4.10 |
| Embryo 9 | 3.18 | 1359 | 228 | 0.040 | 4.30 |
| Embryo 10 | 3.97 | 1413 | 304 | 0.046 | 3.54 |
| Embryo 11 | 3.56 | 1157 | — | — | 3.25 |
| Embryo 12 | 4.56 | 1437 | — | — | 3.15 |

**Supplementary Table S3.** Characteristics of all explant tissues: Untreated explants. For small explants, size is reported as “number of tiles x field of view”; for large explants, absolute area is shown.

| Explant ID | Size | Number of MCCs | Average MCC area ($\mu\text{m}^2$) | Coverage fraction | Density ($\ast 10^{-4}$ MCCs/$\mu\text{m}^2$) |
| --- | --- | --- | --- | --- | --- |
| Explant 1 | $1 \times (680 \times 680) \mu\text{m}^2$ | 33 | 506 | 0.036 | 0.714 |
| Explant 2 | $1 \times (680 \times 680) \mu\text{m}^2$ | 21 | 697 | 0.031 | 0.454 |
| Explant 3 | $2 \times (680 \times 680) \mu\text{m}^2$ | 59 | 840 | 0.053 | 0.638 |
| Explant 4 | $2 \times (680 \times 680) \mu\text{m}^2$ | 81 | 710 | 0.062 | 0.876 |
| Explant 5 | $3 \times (680 \times 680) \mu\text{m}^2$ | 113 | 600 | 0.057 | 0.815 |
| Explant 6 | $2 \times (680 \times 680) \mu\text{m}^2$ | 77 | 497 | 0.041 | 0.833 |
| Explant 7 | $5.18 \text{ mm}^2$ | 262 | 675 | 0.046 | 0.506 |
| Explant 8 | $3.72 \text{ mm}^2$ | 300 | 580 | 0.033 | 0.827 |
| Explant 9 | $7.9 \text{ mm}^2$ | 540 | 735 | 0.043 | 0.683 |

**Supplementary Table S4.** Characteristics of all explant tissues: Treated explants. For small explants, size is reported as “number of tiles × field of view”; for large explants, absolute area is shown.

| Explant ID | Size | Number of MCCs | Average MCC area ($\mu\text{m}^2$) | Coverage fraction | Density ($\ast 10^{-4}$ MCCs/$\mu\text{m}^2$) |
| --- | --- | --- | --- | --- | --- |
| Explant 1 | $1 \times (680 \times 680) \mu\text{m}^2$ | 28 | 536 | 0.032 | 0.606 |
| Explant 2 | $1 \times (680 \times 680) \mu\text{m}^2$ | 16 | 581 | 0.020 | 0.346 |
| Explant 3 | $2 \times (680 \times 680) \mu\text{m}^2$ | 50 | 475 | 0.025 | 0.541 |
| Explant 4 | $1 \times (680 \times 680) \mu\text{m}^2$ | 25 | 540 | 0.029 | 0.540 |
| Explant 5 | $2 \times (680 \times 680) \mu\text{m}^2$ | 56 | 455 | 0.027 | 0.606 |
| Explant 6 | $1.08 \text{ mm}^2$ | 88 | 400 | 0.037 | 0.816 |
| Explant 7 | $1.97 \text{ mm}^2$ | 280 | 405 | 0.056 | 1.412 |
| Explant 8 | $1.75 \text{ mm}^2$ | 194 | 407 | 0.044 | 1.103 |
| Explant 9 | $2.05 \text{ mm}^2$ | 340 | 402 | 0.060 | 1.654 |

**Supplementary Movie S1**

Beating cilia at the surface of an explant, 6 days after explantation. Immobile background has been substracted for a better visualisation of cilia.

**Supplementary Movie S2**

Fluorescent beads transported at the surface of an explan at a distance from the epithelial surface z=20µm.

